# Global mapping of thioredoxin interacting proteins in *Neurospora crassa*

**DOI:** 10.1101/2025.04.24.650474

**Authors:** Lucia Bidondo, Elodie Drula, Jean-Charles Gaillard, Jean Armengaud, Jean-Guy Berrin, Marie-Noëlle Rosso, Lionel Tarrago

**Affiliations:** INRAE, Aix Marseille Univ., BBF, Biodiversité et Biotechnologie Fongiques, Marseille, France; CNRS, Aix Marseille Univ, AFMB, USC1408, Marseille, France; Université Paris-Saclay, CEA, INRAE, Département Médicaments et Technologies pour la Santé (DMTS), SPI, 30200 Bagnols-sur-Cèze, France

**Keywords:** Affinity chromatography, carbohydrate-active enzymes, *Neurospora crassa*, redox regulation, thioredoxin

## Abstract

Thioredoxins (Trx) are essential thiol-oxidoreductases that regulate redox homeostasis by reducing oxidized cysteines in a wide range of target proteins. However, the Trx system and other redox regulation mechanisms remain poorly characterized in saprotrophic filamentous fungi. Here, we identified the components of the *Neurospora crassa* Trx system and uncovered potential redox-regulated proteins using Trx affinity chromatography. Genome search identified three Trx and a single thioredoxin reductase that we named TRX1, TRX2 TRX3 and TRR. Notably, TRX1 carries a C-terminal disordered extension of unknown function, conserved in two ascomycete taxa (*Leotiomycetes* and *Sordariomycetes*). Using recombinant cysteine-to-serine mutants of each Trx, we performed affinity chromatography and identified 1,998 proteins - approximately 19% of the *N. crassa* proteome. To rank the putative Trx targets, we applied a fold enrichment metric, comparing protein abundance before and after affinity chromatography. The average fold enrichment was four, with values reaching up to 117 for the most enriched protein, a DEAD/DEAH box helicase. Among the top-enriched proteins, we identified homologs of known human and plant Trx targets, like peroxiredoxins, as well as 93 transcription factors and 38 kinases. Additional potential Trx targets encompass, ubiquitination-related enzymes, Fe-S cluster assembly proteins, phospholipases, exonucleases, and chitin synthases. Moreover, components of multiprotein complexes were co-purified, reflecting both direct Trx interactions and indirect co-association. Overall, this study provides a global map of potential redox regulated proteins and Trx targets in *N. crassa*, laying the ground for future investigations into redox signaling in filamentous fungi.

## 1. Introduction

Reactive oxygen species (ROS) can regulate metabolic functions and act as signaling molecules, mostly by oxidizing Cys residues into sulfenic acid and disulfide bond inducing alteration of enzyme activity, protein–protein interactions, and subcellular localization [1]. Thioredoxins (Trx) are ubiquitous, small thiol-oxidoreductases capable of reducing oxidized Cys of target proteins, thereby modulating diverse ROS-associated cellular processes. To reduce a target, a Trx uses the two Cys of its conserved WCGPC active site through disulfide exchange [2]. The reduction mechanism is initiated by the nucleophilic attack from N-terminal Cys to the disulfide bond in the target protein, resulting in a transient mixed disulfide formed between both proteins. Subsequently, the C-terminal Cys (also called the “resolving” Cys) reduces the intermolecular bond, releasing the reduced target and the oxidized Trx [3]. The Trx activity can be restored by a Trx reductase (TR), using NADPH as an electron donor [4]. To identify potential Trx targets at proteome scale, an approach was developed taking advantage of the Trx reduction mechanism. By using a Trx mutant in which the resolving Cys of the active site is replaced with a Ser (WCGPS), a stable intermolecular disulfide bond can be formed between the Trx and its targets. The Trx mutant can be immobilized on a column used for affinity chromatography. First, a protein extract of interest is passed through the column, allowing the Trx to trap its targets via disulfide linkage, then after washing, the bound proteins can be eluted using a reducing agent such as dithiothreitol (DTT) and subsequently identified by proteomic analysis [5,6].

Early identified Trx targets include enzymes in which a catalytic Cys requires to be reduced for activity, such as ribonucleotide reductases or methionine sulfoxide reductases [7–9]. Other enzymes were found to be regulated allosterically by disulfide bonds that can be reduced by Trx, like several enzymes involved in photosynthesis [10]. Among Trx targets, several were shown to participate in the regulation of signaling pathways, such as the Yap1 transcription factor in yeast and the apoptosis signal-regulating kinase 1 (ASK1) in mammals, both activated by oxidation and deactivated by Trx-catalyzed reduction [11,12].

While well described in bacteria, yeast, animals, and plants, the knowledge about regulation and signaling associated with ROS and Trx is extremely scarce in filamentous fungi. A wide range of filamentous fungi employs saprotrophic nutrition by decomposing plant cell walls and thereby recycling dead organic matter and playing key roles in global carbon and nitrogen cycling. Composed of structured polysaccharides like cellulose, plant cell walls are highly recalcitrant to degradation, requiring the fungi to secrete a large and diverse set of Carbohydrate-Active enZymes (CAZymes) to breakdown the components into assimilable sugars [13]. The synthesis and secretion of the enzymatic machinery are energetically demanding processes, requiring tight regulation [14]. *Neurospora crassa* is a central model for dissecting the regulatory mechanisms controlling CAZyme expression [15], yet its redox regulatory networks and Trx system remain unexplored.

Animals and bacteria have usually two Trx-encoding genes, while plants genomes can contain large numbers, up to 41 in *Arabidopsis thaliana* [16,17]. *Saccharomyces cerevisiae* genome encodes three Trx: TRX1 and TRX2 localized in the cytoplasm, and a mitochondrial TRX3 which is mitochondrial. It also has two TR, one cytoplasmic (TRR1) and one mitochondrial (TRR2) [18,19]. In the opportunistic pathogen *Aspergillus fumigatus*, one gene encoding a TR was identified and found to be essential for virulence, while in the related species *A. nidulans*, a single cytoplasmic Trx/TR couple was described [20,21]. In the coprophilous fungus *Podospora anserina*, three cytoplasmic Trx were identified [22]. In *N. crassa*, the Trx system was not fully described yet, only one TR-encoding gene (*“Cys-9”*) was identified and its deletion was shown to affect the circadian conidiation rhythm [23].

In this study, we described the Trx system of *N. crassa* and employed Trx-based affinity chromatography combined with deep proteomic analysis to identify potential Trx targets from cellulose-grown cultures. We identified 1,998 proteins and ranked them based on their enrichment fold during affinity purification using a Trx column. Among the top-enriched proteins, we identified homologs of previously established Trx targets, as well as numerous transcription factors and kinases, highlighting both conserved and novel candidates for Trx regulation in *N. crassa*.

## 2. Material and Methods

### 2.1. Identification of the thioredoxin system components in *N. crassa*

The search for the sequences of Trx and TR in *N. crassa* was performed by doing a BLASTP and a BLASTN in *Neurospora crassa* OR74A v2.0 [24] using filtered model proteins and transcripts of the MycoCosm database from the Joint Genome Institute (JGI). The query was based on the thioredoxin system of *Saccharomyces cerevisiae*; TRX1 (Uniprot [25] accession # P22217), TRX2 (P22803), TRX3 (P25372), TRR1 (P29509) and TRR2 (P38816). The predictions for subcellular localization were made using the DeepLoc 2.1 software, standalone version with default parameters [26]. The search for sequences with a C-terminal extension similar to that of the NCU00598 gene product (*N. crassa* TRX1) was conducted using BLASTP against NCBI nr, Uniprot reference proteomes + Swissprot, and MycoCosm databases. The retrieved proteins were aligned using Clustal Omega [27] and manually curated using Jalview 2.11.3.3 [28]. Sequence comparisons of *N. crassa* and *S. cerevisiae* TR and Trx were made using Emboss Needle [29] with default parameters. Structure alignments were performed with Open-Source PyMol 3.1 (https://pymol.org/) using available AlphaFold 2.0 (https://alphafold.ebi.ac.uk/) [30] tridimensional models of *N. crassa* proteins to experimentally structures of *S. cerevisiae* proteins (ScTRR1, PDB#3D8X, [31]; ScTRX1, PDB#3F3Q, [32]; ScTrx2, PDB# 2FA4, [33] and, ScTrx3 PDB# 2OE3, [34]). Prediction of intrinsically disordered region was made using AIUpred (https://aiupred.elte.hu/) [35].

### 2.2. Expression and purification of recombinant thioredoxins

The coding sequences of *N. crassa* TR and Trx isoforms were synthesized with codon optimization for *E. coli* expression and cloned into the pET21(+) expression vector by Twist Bioscience (San Francisco, USA). The cloned TR sequence corresponded to residues 1–334. For the Trx isoforms, both the wild-type and Cys-to-Ser mutant versions were cloned as follows: TRX1 and C35S TRX1 (residues 1–114), TRX2 and C36S TRX2 (residues 1–127), and TRX3 and C74S TRX3 (residues 41–147). All proteins were synthesized with N-terminal His_6_-tag and TEV protease sites except TRX1 and C35S TRX1 that contained the His_6_-tag in C-terminal position. *E. coli* BL21 (DE3) cells were transformed with the expression vectors and grown in Luria-Bertani containing ampicillin (50 μg·ml^−1^) at 37 °C. When the *A*_600_ reached ∼0.5, the production of the recombinant protein was induced by the addition of 100 μM isopropyl β-d-1-thiogalactopyranoside. After 4h incubation at 37 °C, the cells were harvested by centrifugation (5,000 ×g, 15 min) and resuspended in Tris 30 mM pH 8.0, 0.5 M NaCl and 25 mM imidazole, containing Pierce protease inhibitor (ThermoFisher Scientific) and frozen at – 20 °C until protein purification. After thawing, cellular lysis was performed in ice by sonication (1 min, pulse on 10 s, pulse off 10 s and amplitude 30%, repeated twice) using a 505 Sonicator (Fisher Scientific). The soluble proteins were separated from cell debris by centrifugation for 1 h at 4 °C at 15,000 ×g. Proteins were purified on Ni-columns (HisTrap FF crude, Cytiva) in Tris 30 mM pH 8.0, 0.5 M NaCl and 500 mM imidazole. Then, the proteins were concentrated by ultrafiltration (Vivaspin™ Centrifugal Concentrator, 10kDa, Cytivia) and desalted in Tris 30 mM pH 8.0 using PD10 columns (Cytivia). For further purification, TRR was diluted 10 times in Tris 30 mM pH 7.0, loaded in anion exchange column (Resource Q, Cytiva), and eluted with a linear gradient of salt to 0.5M using Tris 30 mM 1 M NaCl pH 7.0. TRX1, C35S TRX1 and C74S TRX3 were purified similarly by cation exchange on Resource S column (Cytiva). TRX1 and C74S TRX3 were further purified by gel filtration (HiLoad^TM^ 16/1600 Superdex^TM^, Sigma-Aldrich®) in Tris 30 mM pH 8.0. Each chromatography was performed using the AKTA Pure system (Cytiva). The fractions were analyzed by SDS-PAGE using Bolt Bis-Tris 10% or 12% acrylamide gels in MES-SDS running buffer (ThermoFisher Scientific) and stained using SafeStain SimplyBlue (ThermoFisher Scientific). The fractions for which the protein was apparently pure were pooled and the protein solution was concentrated by ultrafiltration. The protein final concentration was determined by absorbance at 280 nm using the theorical extinction coefficient calculated using the Expasy ProtParam tool [36].

### 2.3. Turbidimetric assay of insulin disulfide reduction

The activity of the wild-type Trx and Cys-to-Ser Trx mutant was measured as described [37] with a few modifications. Briefly, 5 μM of Trx was incubated in 0.1 M K_2_HPO_4_, 2 mM EDTA and pH 7.5 with 1 mM DTT. After 15 min, 130 μM of bovine pancreatic insulin (Sigma-Aldrich®) was added and the precipitation of insulin chain B was followed by the absorbance at 650 nm at 23 °C for 45 minutes.

### 2.4. Extraction of mycelium soluble proteins

*Neurospora crassa* WT (FGSC#2489) was grown in 250 mL baffled Erlenmeyer flask containing 100 mL of Vogel’s minimal medium (VMM) [38] and inoculated with 1×10^6^ conidia.mL^−1^. The cultures were incubated for 72 h at 25 °C with shaking at 115 rpm in Multitron Standard incubator (Infors HT). The mycelium was filtered in Miracloth, washed with 0.9% NaCl, flash-frozen with liquid nitrogen and stored at −80 °C. The mycelium was cryomilled and 15 mL of extraction buffer (0.1 M K_2_HPO_4_, 2 mM Ethylenediamine tetraacetic acid (EDTA), 250 mM NaCl, 1 mM phenylmethylsulfonyl fluoride (PMSF) pH 7.5) was added. The extraction mixture was incubated for 1.5 h at room temperature under gentle shaking. The soluble protein fraction was separated by centrifugation at 15,000×g at 4°C for 1 h and the supernatant was filtered (0.45 μm). The protein extracts from four biological replicates were pooled prior to affinity chromatography. The protein concentration was determined using Bradford assay (Bio-Rad Protein Assay Dye Reagent Concentrate).

### 2.5. Immobilization of Cys-to-Ser mutated Trx and affinity chromatography

The C35S TRX1, C36S TRX2 or C74S TRX3 (10 mg) were immobilized on 1 mL HiTrap® NHS-activated Sepharose High Performance (Cytiva) following the supplier’s recommendations. Briefly, the column was activated with 1 mM of ice-cold HCl. Immediately, 1 mL of Trx solution (10 mg.mL^−1^) was applied and incubated for 30 min at 25 °C. Then, the column was washed with three column volumes (CV) of coupling buffer (200 mM NaHCO_3_, 500 mM NaCl, pH 8.3). Free activated *N*-hydroxysuccinimide sites were blocked with 500 mM ethanolamine, 500 mM NaCl, pH 8.3. HiTrap Trx-Sepharose column was connected to AKTA pure system (Cytiva). The immobilized Trx was reduced with 3 CV of elution buffer (0.1 M K_2_HPO_4_, 2 mM EDTA, 250 mM NaCl, 2 mM DTT, pH 7.5) at 1 mL.min^−1^ and equilibrated with 10 CV of binding buffer (0.1 M K_2_HPO_4_, 2 mM EDTA, 250 mM NaCl, pH 7.5) at 1 mL.min^−1^. Total soluble proteins of *N. crassa* (100 mg or 10 mg) were loaded onto the column at a flow rate of 0.7 mL.min^−1^. The column was washed with 20 CV of binding buffer and the retained proteins were eluted with 10 CV of elution buffer at 1 mL.min^−1^ providing the eluates *“E_100mg_” and “E_10mg_”*, respectively. The eluted fractions were concentrated in 3 kDa Vivaspin™ ultrafiltration spin columns (Cytiva) and the protein concentration was determined using Bradford assay. Intracellular proteins and of each *“E_100mg_” and “E_10mg_”* were identified after trypsin digestion of 10 µg and liquid chromatography coupled to high resolution tandem mass spectrometry (LC-MS/MS) performed with an Exploris 480 (Thermo) tandem mass spectrometer as described [39]. The data interpretation was performed similarly to [40].

### 2.6. Identification of homologs of human Trx targets

To identify *N. crassa* homologs of human Trx targets, the sequences of 341 manually curated human TRX1 and TRX2 targets [41] were retrieved from UniProt in FASTA format. Using standalone BLAST+ software, the human targets were aligned against the *N. crassa* proteome (Uniprot accession UP000001805). The candidate homologs were selected based on the lowest E-value and highest alignment percentage, with the thresholds E-value < 1e^−10^, alignment length ≥ 60%. The annotations and EC numbers of the retrieved *N. crassa* proteins were manually inspected to exclude doubtful proteins.

## 3. Results

### 3.1. The *N. crassa* thioredoxin system is composed of one TR and three Trx

To identify the genes encoding TR and Trx in *N. crassa*, we performed BLAST searches using the *S. cerevisiae* TR and Trx protein sequences as query. We found only one gene (NCU08352, c*ys-9*) encoding a potential TR, with a predicted molecular weight of ∼35.9 KDa (**Table 1**). Protein sequence alignment revealed high identity with the yeast homologs (65.8% with TRR1 and 59.6% with TRR2) (**Figure S1**). Structural superimposition of the predicted *N. crassa* TRR model with the crystallographic structure of yeast TRR1 confirmed strong conservation of the overall fold. Moreover, *N. crassa* TRR contains all the characteristic residues of a low-molecular-weight dimeric thioredoxin reductase, similar to its yeast counterparts [42]. It has no N-terminal peptide signal, but a C-terminal extension of undetermined function, predicted to be disordered. The protein is predicted to be localized in the cytoplasm and/or the peroxisome (**Table 1**). Using the same approach, we found three Trx-encoding genes (NCU00598, NCU05731, NCU06556). We named the proteins TRX1, TRX2, and TRX3 based on their chromosomal locations and their predicted subcellular localization in analogy with the *S. cerevisiae* Trx. The three Trx share ∼25 to ∼52 % identity with the *S. cerevisiae* protein sequences and a very well-conserved thioredoxin fold with a typical WCGPC active site (**Table 1**, **Figure S2**). Interestingly, the gene NCU00598 encodes for two potential isoforms of TRX1, both having a C-terminal undetermined function, predicted to be disordered (**Table 1**, **Figure S2, S3**). They differ in their C-terminal extension sequences, with one variant having 21 extra amino acids due to alternative splicing that retains the second intron (**Figure S2, S3**). Available RNAseq data [43] indicate that only the shortest transcript with removal of the second intron is expressed (**Figure S3**). To evaluate if similar C-terminal extension can be found in other organisms, we performed BLAST searches and identified 151 sequences with a similar extension, only from other Ascomycetes fungi of *Leotiomycetes* and *Sordariomycetes* classes (**Table S1**). Notably, all these sequences share a C-terminal extension like the one resulting from the translation of the shortest transcript of NCU00598, suggesting that the longest transcript might result from missannotation or be produced only under specific conditions (**Figure S3**). The TRX1 resulting from the NCU00598 gene expression in which the two introns are spliced has a theoretical molecular weight of ∼22.8 kDa and is predicted to be localized in cytoplasm and/or the endoplasmic reticulum (**Table 1**). The genes NCU05731 and NCU06556 code for the putative TRX2 of ∼13.7 kDa predicted to be localized in the cytoplasm, and the potential TRX3 of ∼16.1 kDa having a N-terminal extension of 35 amino acids predicted to be a mitochondrial signal peptide, respectively (**Table 1**, **Figure S2**). To test their ability to reduce disulfide bonds, we produced and purified the Trx, without the signal peptide for TRX3, and in the case of TRX1, we removed the C-terminal extension after unsuccessful attempts to obtain the full-length protein. The three enzyme efficiently reduced disulfide bonds in insulin (**Figure S4**).

**Table 1.**
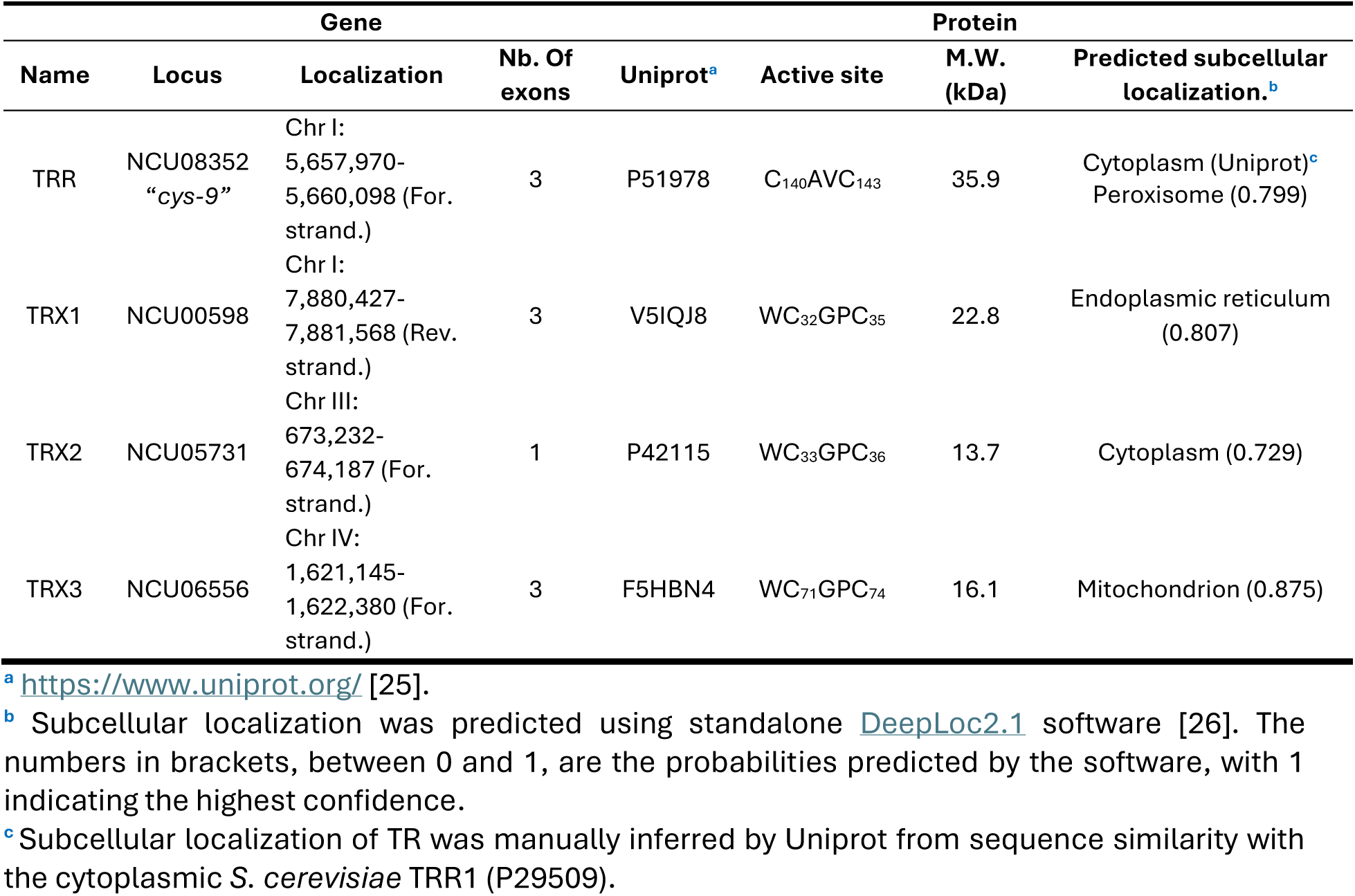
Description of *N. crassa* TR and Trx genes and protein sequences.

Altogether, these results showed that the Trx system in *N. crassa* resembles that of the baker yeast, although it seems to lack a mitochondrial TR.

### 3.2. Fold enrichment serves as a metric to rank candidate Trx targets

To identify Trx targets in *N. crassa*, we performed affinity chromatography with Cys-to-Ser mutants of TRX1, TRX2 and TRX3, independently. Our hypothesis was that each Trx may exhibit distinct target specificities, potentially reflecting differences in subcellular localization or redox interactions. The recombinant C35S TRX1, C36S TRX2, and C74S TRX3 proteins were produced similarly to their wild-type counterparts and, we confirmed their loss of disulfide-reducing activity (**Figure S4**). Then, 10 mg of each Trx mutant was bound to a sepharose column for affinity chromatography (**Figure 1A**). To induce saprotrophic metabolism and CAZyme secretion [44], *N. crassa* was grown on cellulose for three days, then soluble proteins from four biological replicates were extracted and pooled, then used for affinity chromatography. For each Trx isoform, we applied on the column two different amounts of *N. crassa* proteins: 100 mg, as a saturating quantity, and 10 mg, equivalent to the amount of Trx immobilized on the column. Our hypothesis was that the retention of the targets on the column depends not only on its abundance in the initial extract, but also on its affinity for the Trx. Under saturating conditions, abundant proteins are more likely to be preferentially retained, whereas under limiting conditions, proteins with high affinity but lower abundance may be more efficiently captured. Following affinity chromatography, we obtained two eluates for each Trx isoform, referred to as E*_100mg_* and E*_10mg_*, resulting in a total of six samples (**Figure 1A**). The proteins in each eluate were identified by LC-MS/MS. A sample of the initial protein extract was analyzed in parallel and used as a reference proteome. Across the seven samples, we identified a total of 2,546 proteins (**Data Set 1**). Among these, 1,998 unique proteins were detected in at least one of the affinity chromatography eluates, while 548 were found exclusively in the reference proteome and are therefore unlikely to be Trx targets (**Table 2**, **Data Set 1**). The number of proteins identified was comparable across the six affinity chromatography samples, ranging from 1,359 in TRX3 E*_10mg_* to 1,716 in TRX1 E*_10mg_* (**Table 2**). To prioritize candidate Trx targets, we calculated a fold enrichment value for each protein, based on the assumption that true interactors would be retained in the eluates while low-affinity proteins would be lost during washing. The fold enrichment was defined as the ratio of a protein’s relative abundance in the eluate to its relative abundance in the reference proteome (**Table 2**). Across the six eluates, the mean fold enrichment varied from 4.0 in TRX1 E*_100mg_* to 5.1 in TRX3 E*_10mg_*, while the highest fold enrichment reached 117 (TRX3 E*_10mg_*). In each case, approximately one-third of the proteins displayed a fold enrichment above or equal to four, indicating strong retention on the column and suggesting specific interaction with the immobilized Trx mutant. Around one third of proteins showed fold enrichment between one and four, meaning their abundance was higher or equal to those in the reference proteome. And finally, approximately one-third of the proteins showed a fold enrichment value below one, indicating lower abundance after affinity chromatography compared to the reference proteome. This suggests that these proteins likely did not interact with Trx via disulfide bond. Similarly, several proteins identified lacked Cys residues and were potentially retained nonspecifically due to high abundance in the input sample, as previously suggested [27]. To explore this further, we compared the fold enrichment of proteins containing Cys to those without Cys across the six eluates (**Figure 1B**). Proteins with Cys showed fold enrichment values ranging from 4.1 (in TRX1 E*_100mg_*) to 5.2 (in TRX3 E*_10mg_*). In contrast, proteins lacking Cys exhibited significantly lower enrichment, with values ranging from 1.5 (in TRX3 E*_100mg_*) to 2.1 (in TRX3 E*_10mg_*). In each eluate, the mean of fold enrichment of protein without Cys was more than two-fold lower than the mean of fold enrichment of Cys-containing proteins. As expected, proteins with Cys residues exhibited significantly higher fold enrichment compared to proteins lacking Cys, validating the use of fold enrichment as a reliable indicator of specific Trx interactions. Consequently, proteins with the highest fold enrichment are more likely to be promising Trx targets candidates in *N. crassa*.

**Figure 1.**
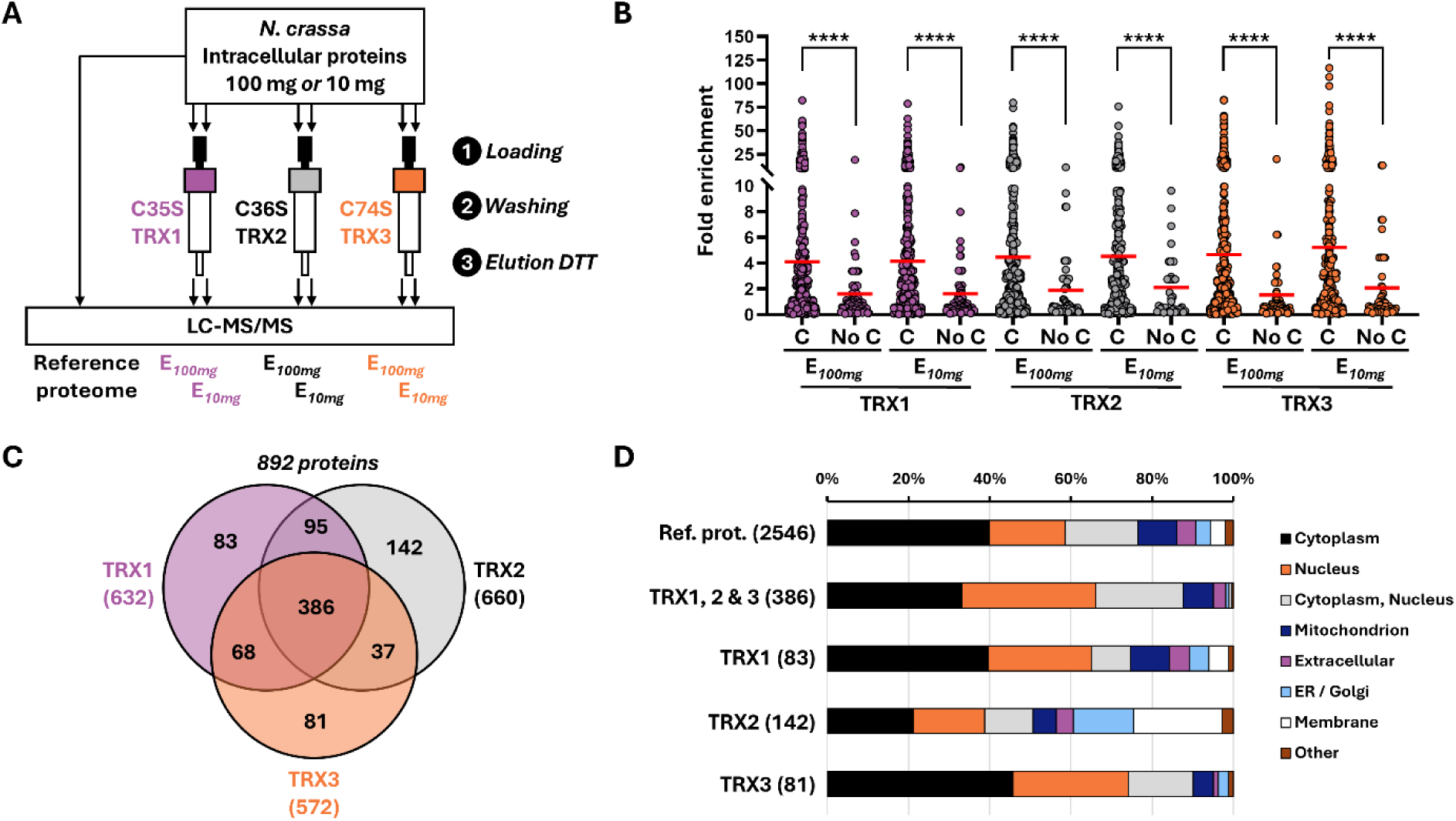
Identification of Trx-interacting proteins by affinity chromatography with fold enrichment calculation. **A)** *N. crassa* was cultured with 2% cellulose, and intracellular proteins were extracted. Affinity chromatography was performed for each Trx using 100 mg or 10 mg of extracted proteins loaded onto columns containing C35S TRX1, C36S TRX2, or C74S TRX3 (*Step 1*). After washing (*Step 2*), the retained proteins were eluted with DTT (*Step 3*), yielding two sets of eluted proteins per Trx: “E*_100mg_*” and “E*_100mg_*”. Protein identity and abundance in the six elution samples and the initial extract (“Reference proteome”) were determined by LC-MS/MS. The “fold enrichment” was calculated by dividing the relative abundance of each protein in the elution by its abundance in the reference proteome. **B)** Fold enrichment comparison of proteins with at least one Cys (“C”) versus those without Cys (“No C”) in E*_100mg_* and E*_10mg_* for each Trx. Each dot represents a protein, with the mean fold enrichment indicated by a horizontal *red* bar. Statistical analysis was performed using an unpaired Mann-Whitney t-test (\*\*\*\**P* < 0.0001). **C)** Venn diagram of proteins found in both E*_100mg_* and E*_10mg_* for each Trx with a fold enrichment ≥ 4. **D)** Proportions of the subcellular localizations of proteins with a fold enrichment ≥ 4 that were common among all eluted proteins (*“TRX1, 2 & 3”*), or unique to TRX1, TRX2, or TRX3 eluates (*“TRX1”*, *“TRX2”* and *“TRX3”*, respectively). These groups correspond to those of the Venn diagram (Panel C). Proportions were compared to those found in the reference proteome (*“Ref. prot.”*). The number of proteins is indicated in brackets.

**Table 2.** Fold enrichment of proteins found in E*_100mg_* and E*_10mg_* of each Trx affinity chromatography.

|  | TRX1 |  | TRX2 |  | TRX3 |  | Total |
| --- | --- | --- | --- | --- | --- | --- | --- |
| | $E_{100mg}$ | $E_{10mg}$ | $E_{100mg}$ | $E_{10mg}$ | $E_{100mg}$ | $E_{10mg}$ | |
| Number of identified proteins | 1697 | 1716 | 1467 | 1448 | 1454 | 1359 | 1998 |
| Mean fold enrichment for all proteins | 4.0 | 4.1 | 4.4 | 4.5 | 4.6 | 5.1 | 4.5 |
| Max fold enrichment | 82 | 79 | 80 | 76 | 83 | 117 | 117 |
| Nb. of prot. with fold enrichment $\geq 4$ | 481 | 511 | 495 | 559 | 432 | 481 | 892 |
| Nb. of prot. with $1 < \text{fold enrichment} < 4$ | 649 | 660 | 491 | 436 | 527 | 419 | 576 |
| Nb. of prot. with fold enrichment $< 1$ | 567 | 545 | 481 | 453 | 495 | 459 | 530 |

### 3.3. Homologs of known Trx targets were enriched by affinity chromatography

Among the proteins retained by Trx during affinity chromatography, we identified homologs of known Trx targets. Most of them were retained by the three Trx and we present only their highest fold enrichment value (**Data Set 1**). These proteins include methionine sulfoxide reductases A and B (NCU10029, NCU03891) [8,9] and a mitochondrial peroxiredoxin (NCU06031) [45], which exhibited maximum fold enrichments of 3.7, 1.9, and 5.4, respectively, in TRX3 E*_100mg_*. We also found an AP1/BZIP transcription factor (NCU03905), homologous to *S. cerevisiae* Yap1 [11,46], with a maximum fold enrichment of 5.6 in TRX2 E*_100mg_*. Additionally, a PAPS reductase (NCU02005) [47] and the large chain of the ribonucleoside-diphosphate reductase (NCU08514) [7] were identified with maximal fold enrichments of 24.9, and 15.7 in TRX3 E*_10mg_* and TRX3 E*_100mg_*, respectively (**Data Set 1**). Moreover, a peptidyl-prolyl isomerase (NCU08514), for which a homolog was shown to be activated by Trx in plants [48], was present among the top 50 most enriched proteins, with a maximal fold enrichment of 36.9 in TRX3 E*_10mg_* (**Data Set 1**, **Table 3**). To compare our results with previous findings in a more systematic approach, we searched for *N. crassa* homologs in a curated list of 341 human TRX1 and TRX2 targets [41] and found that 82 were identified in our study (**Data Set 2**). Of these, 53 had fold enrichment values between one and four and 24 exceeding four. The mean fold enrichment was ∼3.6, with the highest value of 98 observed for the catalytic subunit PAN2 of the deadenylation complex PAN2-PAN3, which was the third most enriched protein in all affinity chromatography eluates (**Table 3**). Consistent with previous studies [5,6], these results confirm that affinity chromatography effectively identifies relevant Trx targets and supports the use of fold enrichment values as a reliable method for Trx targets identification.

**Table 3.**
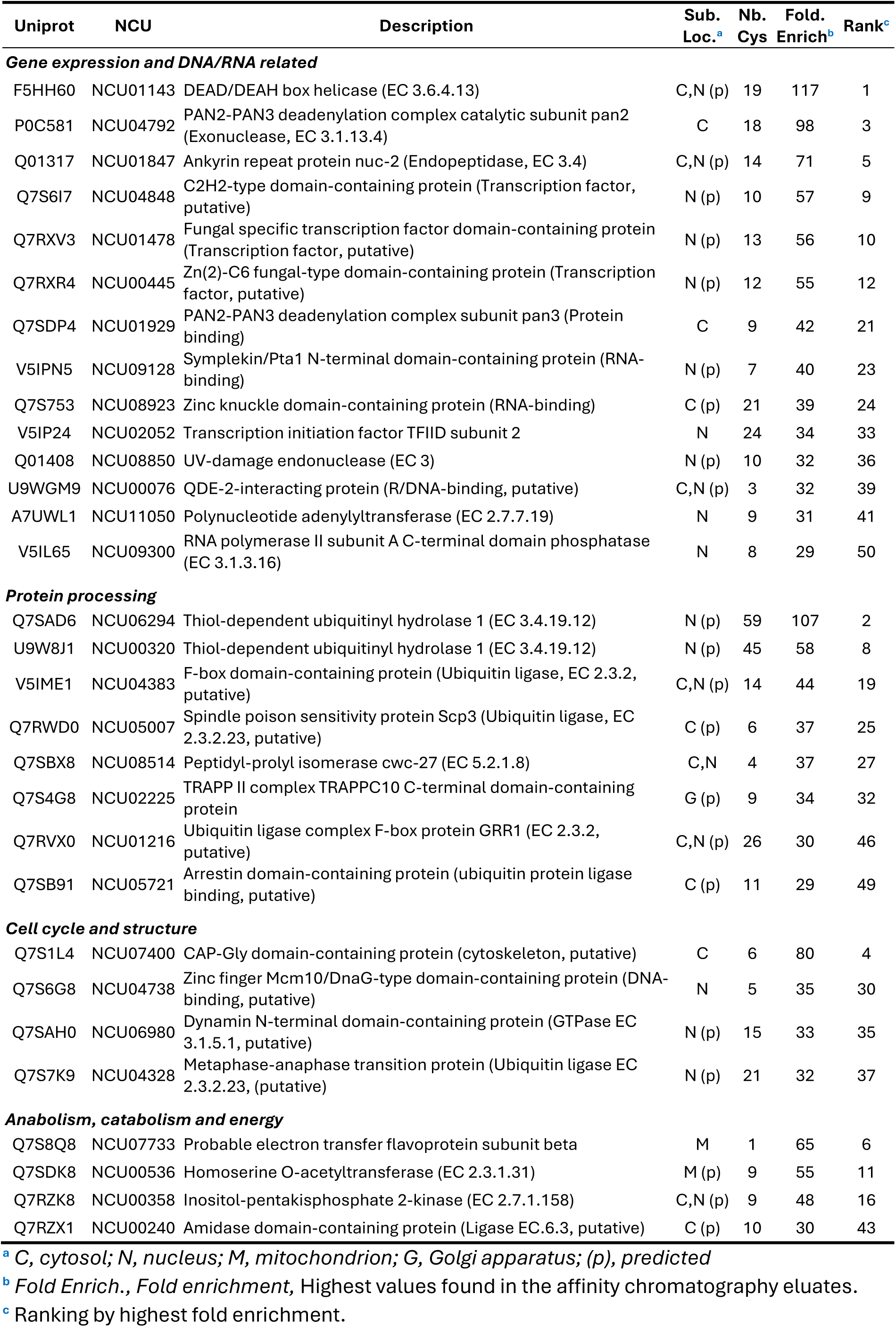
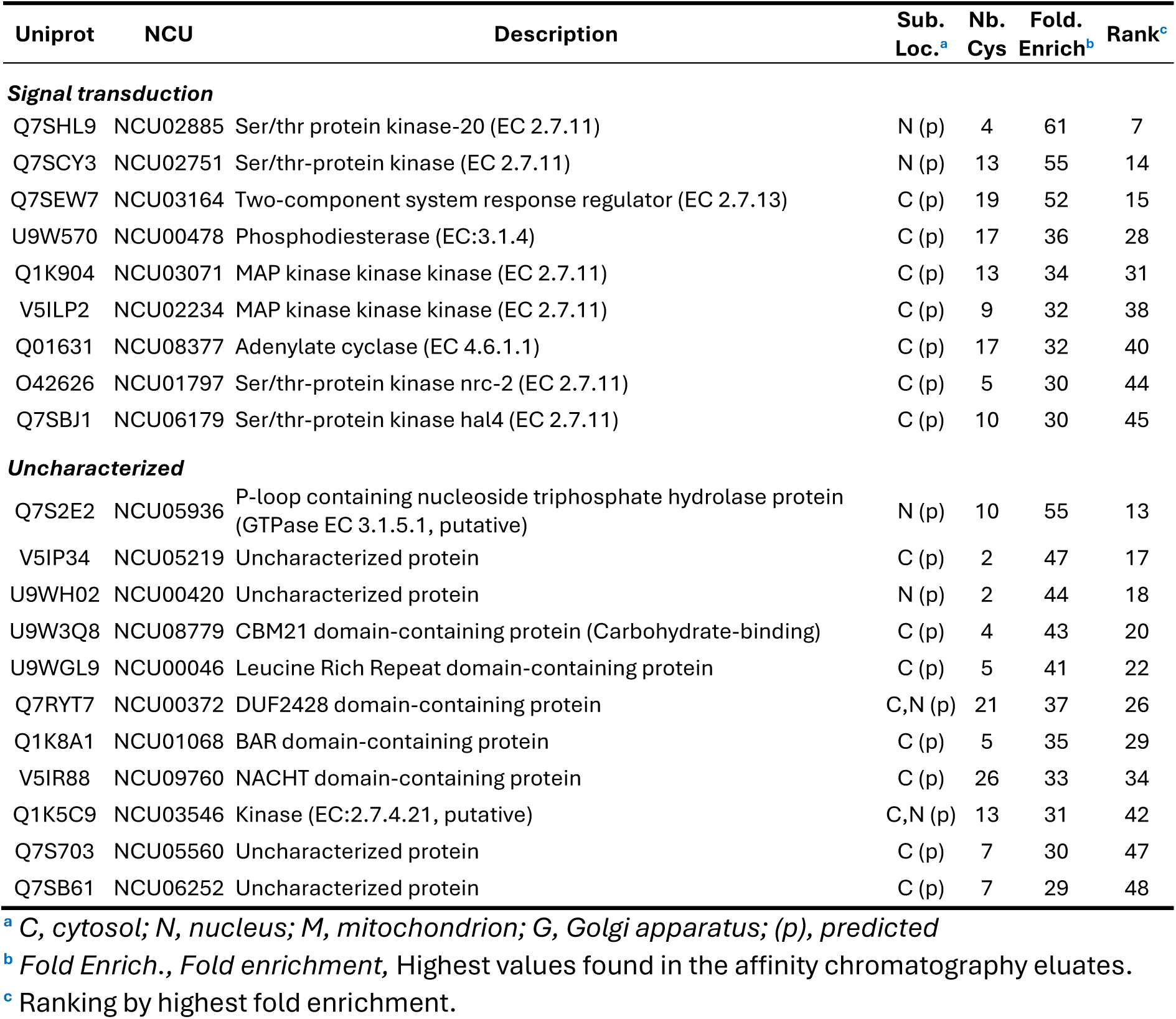
Top 50 enriched proteins in Trx affinity chromatography.

| Uniprot | NCU | Description | Sub. Loc.<sup>a</sup> | Nb. Cys | Fold. Enrich<sup>b</sup> | Rank<sup>c</sup> |
| --- | --- | --- | --- | --- | --- | --- |
| **Gene expression and DNA/RNA related** |  |  |  |  |  |  |
| F5HH60 | NCU01143 | DEAD/DEAH box helicase (EC 3.6.4.13) | C,N (p) | 19 | 117 | 1 |
| P0C581 | NCU04792 | PAN2-PAN3 deadenylation complex catalytic subunit pan2 (Exonuclease, EC 3.1.13.4) | C | 18 | 98 | 3 |
| Q01317 | NCU01847 | Ankyrin repeat protein nuc-2 (Endopeptidase, EC 3.4) | C,N (p) | 14 | 71 | 5 |
| Q7S6I7 | NCU04848 | C2H2-type domain-containing protein (Transcription factor, putative) | N (p) | 10 | 57 | 9 |
| Q7RXV3 | NCU01478 | Fungal specific transcription factor domain-containing protein (Transcription factor, putative) | N (p) | 13 | 56 | 10 |
| Q7RXR4 | NCU00445 | Zn(2)-C6 fungal-type domain-containing protein (Transcription factor, putative) | N (p) | 12 | 55 | 12 |
| Q7SDP4 | NCU01929 | PAN2-PAN3 deadenylation complex subunit pan3 (Protein binding) | C | 9 | 42 | 21 |
| V5IPN5 | NCU09128 | Symplekin/Pta1 N-terminal domain-containing protein (RNA-binding) | N (p) | 7 | 40 | 23 |
| Q7S753 | NCU08923 | Zinc knuckle domain-containing protein (RNA-binding) | C (p) | 21 | 39 | 24 |
| V5IP24 | NCU02052 | Transcription initiation factor TFIID subunit 2 | N | 24 | 34 | 33 |
| Q01408 | NCU08850 | UV-damage endonuclease (EC 3) | N (p) | 10 | 32 | 36 |
| U9WGM9 | NCU00076 | QDE-2-interacting protein (R/DNA-binding, putative) | C,N (p) | 3 | 32 | 39 |
| A7UWL1 | NCU11050 | Polynucleotide adenyltransferase (EC 2.7.7.19) | N | 9 | 31 | 41 |
| V5IL65 | NCU09300 | RNA polymerase II subunit A C-terminal domain phosphatase (EC 3.1.3.16) | N | 8 | 29 | 50 |
| **Protein processing** |  |  |  |  |  |  |
| Q7SAD6 | NCU06294 | Thiol-dependent ubiquitinyl hydrolase 1 (EC 3.4.19.12) | N (p) | 59 | 107 | 2 |
| U9W8J1 | NCU00320 | Thiol-dependent ubiquitinyl hydrolase 1 (EC 3.4.19.12) | N (p) | 45 | 58 | 8 |
| V5IME1 | NCU04383 | F-box domain-containing protein (Ubiquitin ligase, EC 2.3.2, putative) | C,N (p) | 14 | 44 | 19 |
| Q7RWD0 | NCU05007 | Spindle poison sensitivity protein Scp3 (Ubiquitin ligase, EC 2.3.2.23, putative) | C (p) | 6 | 37 | 25 |
| Q7SBX8 | NCU08514 | Peptidyl-prolyl isomerase cwc-27 (EC 5.2.1.8) | C,N | 4 | 37 | 27 |
| Q7S4G8 | NCU02225 | TRAPP II complex TRAPPC10 C-terminal domain-containing protein | G (p) | 9 | 34 | 32 |
| Q7RVX0 | NCU01216 | Ubiquitin ligase complex F-box protein GRR1 (EC 2.3.2, putative) | C,N (p) | 26 | 30 | 46 |
| Q7SB91 | NCU05721 | Arrestin domain-containing protein (ubiquitin protein ligase binding, putative) | C (p) | 11 | 29 | 49 |
| **Cell cycle and structure** |  |  |  |  |  |  |
| Q7S1L4 | NCU07400 | CAP-Gly domain-containing protein (cytoskeleton, putative) | C | 6 | 80 | 4 |
| Q7S6G8 | NCU04738 | Zinc finger Mcm10/DnaG-type domain-containing protein (DNA-binding, putative) | N | 5 | 35 | 30 |
| Q7SAH0 | NCU06980 | Dynamin N-terminal domain-containing protein (GTPase EC 3.1.5.1, putative) | N (p) | 15 | 33 | 35 |
| Q7S7K9 | NCU04328 | Metaphase-anaphase transition protein (Ubiquitin ligase EC 2.3.2.23, putative) | N (p) | 21 | 32 | 37 |
| **Anabolism, catabolism and energy** |  |  |  |  |  |  |
| Q7S8Q8 | NCU07733 | Probable electron transfer flavoprotein subunit beta | M | 1 | 65 | 6 |
| Q7SDK8 | NCU00536 | Homoserine O-acetyltransferase (EC 2.3.1.31) | M (p) | 9 | 55 | 11 |
| Q7RZK8 | NCU00358 | Inositol-pentakisphosphate 2-kinase (EC 2.7.1.158) | C,N (p) | 9 | 48 | 16 |
| Q7RZX1 | NCU00240 | Amidase domain-containing protein (Ligase EC.6.3, putative) | C (p) | 10 | 30 | 43 |

<sup>a</sup> C, cytosol; N, nucleus; M, mitochondrion; G, Golgi apparatus; (p), predicted<sup>b</sup> Fold Enrich., Fold enrichment, Highest values found in the affinity chromatography eluates.<sup>c</sup> Ranking by highest fold enrichment.

**Table 3 (continued). Top 50 enriched proteins in Trx affinity chromatography**
| Uniprot | NCU | Description | Sub. Loc.<sup>a</sup> | Nb. Cys | Fold. Enrich<sup>b</sup> | Rank<sup>c</sup> |
| --- | --- | --- | --- | --- | --- | --- |
| **Signal transduction** |  |  |  |  |  |  |
| Q7SHL9 | NCU02885 | Ser/thr protein kinase-20 (EC 2.7.11) | N (p) | 4 | 61 | 7 |
| Q7SCY3 | NCU02751 | Ser/thr-protein kinase (EC 2.7.11) | N (p) | 13 | 55 | 14 |
| Q7SEW7 | NCU03164 | Two-component system response regulator (EC 2.7.13) | C (p) | 19 | 52 | 15 |
| U9W570 | NCU00478 | Phosphodiesterase (EC:3.1.4) | C (p) | 17 | 36 | 28 |
| Q1K904 | NCU03071 | MAP kinase kinase kinase (EC 2.7.11) | C (p) | 13 | 34 | 31 |
| V5ILP2 | NCU02234 | MAP kinase kinase kinase (EC 2.7.11) | C (p) | 9 | 32 | 38 |
| Q01631 | NCU08377 | Adenylate cyclase (EC 4.6.1.1) | C (p) | 17 | 32 | 40 |
| O42626 | NCU01797 | Ser/thr-protein kinase nrc-2 (EC 2.7.11) | C (p) | 5 | 30 | 44 |
| Q7SBJ1 | NCU06179 | Ser/thr-protein kinase hal4 (EC 2.7.11) | C (p) | 10 | 30 | 45 |
| **Uncharacterized** |  |  |  |  |  |  |
| Q7S2E2 | NCU05936 | P-loop containing nucleoside triphosphate hydrolase protein (GTPase EC 3.1.5.1, putative) | N (p) | 10 | 55 | 13 |
| V5IP34 | NCU05219 | Uncharacterized protein | C (p) | 2 | 47 | 17 |
| U9WH02 | NCU00420 | Uncharacterized protein | N (p) | 2 | 44 | 18 |
| U9W3Q8 | NCU08779 | CBM21 domain-containing protein (Carbohydrate-binding) | C (p) | 4 | 43 | 20 |
| U9WGL9 | NCU00046 | Leucine Rich Repeat domain-containing protein | C (p) | 5 | 41 | 22 |
| Q7RYT7 | NCU00372 | DUF2428 domain-containing protein | C,N (p) | 21 | 37 | 26 |
| Q1K8A1 | NCU01068 | BAR domain-containing protein | C (p) | 5 | 35 | 29 |
| V5IR88 | NCU09760 | NACHT domain-containing protein | C (p) | 26 | 33 | 34 |
| Q1K5C9 | NCU03546 | Kinase (EC:2.7.4.21, putative) | C,N (p) | 13 | 31 | 42 |
| Q7S703 | NCU05560 | Uncharacterized protein | C (p) | 7 | 30 | 47 |
| Q7SB61 | NCU06252 | Uncharacterized protein | C (p) | 7 | 29 | 48 |

<sup>a</sup> C, cytosol; N, nucleus; M, mitochondrion; G, Golgi apparatus; (p), predicted
<sup>b</sup> Fold Enrich., Fold enrichment, Highest values found in the affinity chromatography eluates.
<sup>c</sup> Ranking by highest fold enrichment.

### 3.4. The initial quantity of protein does not affect the outcome of affinity chromatography

To assess whether the quantity of loaded proteins influenced the outcome of the affinity chromatography, we compared both the identity and fold enrichment of proteins detected in the E*_100mg_* and E*_10mg_* eluates for each Trx isoform (**Figure S5**). For TRX1, 1,880 proteins were identified in total, with 1,533 (∼82%) shared between both conditions, while 164 (∼9%) and 183 (∼10%) were specific to E*_100mg_* and E*_10mg_*, respectively. Similarly, TRX2 yielded 1,644 proteins, of which 1,271 (∼77%) were common, and 196 (∼12%) and 177 (∼11%) were unique to the high and low quantity of loaded proteins, respectively. For TRX3, 1,600 proteins were detected overall, with 1,213 (∼76%) found in both eluates, and 241 (∼15%) and 146 (∼9%) unique to E*_100mg_* and E*_10mg_*, respectively. These results indicate a strong overlap between the two quantities of protein, suggesting that the amount of extract loaded onto the column has only a minor influence on the overall identity of proteins retained by each Trx isoform. Further analysis of the fold enrichment values confirmed this observation. For TRX1 and TRX2, there was no significant difference in fold enrichment of the proteins identified in both eluates, while the difference was only slightly higher for TRX3 E*_10mg_* (**Figure S5**). When plotting the fold enrichment in E*_10mg_* as a function of that in E*_100mg_*, we observed positive correlations for all Trx isoforms, with regression slopes between 0.8 and 1.0 and R² values exceeding 0.7 (**Figure S5**), further supporting the consistency between both conditions. Altogether, these results demonstrate that using either high (100 mg) or low (10 mg) quantity of protein for affinity chromatography leads to largely comparable results, both in terms of protein identity and fold enrichment, in our conditions. Therefore, in subsequent analyses, we merged the data from both E*_100mg_* and E*_10mg_* eluates for each Trx isoform and retained the highest fold enrichment value for each individual protein.

### 3.5. Nuclear proteins are overrepresented among *N. crassa* Trx candidate targets

To obtain a more precise view of the potential Trx targets in *N. crassa*, we focused our analysis on proteins with a fold enrichment equal to or greater than four. A total of 892 unique proteins met this criterion with 632, 660, and 572 proteins detected in TRX1, TRX2, and TRX3 affinity chromatography, respectively (**Figure 1C**). Among these, 386 proteins (43%) were common to all three Trx isoforms, while 83 (22%), 142 (37%), and 81 (4%) were uniquely enriched in TRX1, TRX2, and TRX3 eluates, respectively. We next compared the mean fold enrichment of proteins common to all three Trx to that of proteins uniquely associated with a single Trx. The common proteins exhibited a mean fold enrichment of approximately 18, whereas the isoform-specific proteins showed markedly lower values, around six for each group (**Figure S6**). Taken together, these results indicate that, under the current experimental conditions, the three *N. crassa* Trx display limited specificity in distinguishing their targets by affinity chromatography. However, since the three Trx isoforms are potentially targeted to distinct subcellular compartments, namely the endoplasmic reticulum, cytosol, and mitochondrion for TRX1, TRX2, and TRX3, respectively, we investigated whether the proteins uniquely enriched by each isoform were localized to the corresponding compartment. To this end, we first compiled the subcellular localization of all the identified proteins using both prediction and Uniprot annotation (**Data Set 1**). Then, we compared the distribution of subcellular localizations among proteins uniquely associated with each Trx isoform, those shared by all three isoforms, and the entire *N. crassa* reference proteome (**Figure 1D**). In the reference proteome, most of the 2,546 proteins were localized in the cytoplasm (39.8%) and the nucleus (18.7%), or addressed to both compartments (18.0%). Mitochondrial proteins accounted for 9.5%, while extracellular, ER/Golgi, and membrane-associated proteins represented 4.7%, 3.7%, and 3.6%, respectively. The remaining 2.0% were assigned to other compartments. For the 83 proteins found exclusively in TRX1 affinity chromatography, the proportion of nuclear proteins increased to 25% while the one of endoplasmic reticulum (4.8%), and the other localizations, were barely different from the reference proteome. For the 142 proteins found only in TRX2 column, the proportion of cytoplasmic proteins decreased strongly, to 21.1% while those of ER/Golgi and membrane associated proteins increased three and five fold (11.1% and 18.2% respectively). In the case of the 81 proteins found only in TRX3 affinity chromatography, the proportion of cytoplasmic and nuclear proteins increased to 45.7% and 28.4%, respectively, the proportion of mitochondrial proteins decreased to 4.9%. These observations indicate that the proteins uniquely enriched by each Trx do not predominantly localize to their expected subcellular compartments. In particular, the lack of enrichment for mitochondrial proteins with TRX3 and the unexpected increase in ER/Golgi and membrane-associated proteins among TRX2 candidate targets showed that, in our experimental conditions, the three *N. crassa* Trx had limited selectivity toward compartment-specific putative targets (**Figure 1D**). Next, we performed the same analysis for the 386 proteins commonly found among the three Trx (**Figure 1D**). Strikingly, the proportions of nuclear and cytoplasmic/nuclear proteins increased to 32.9% and 21.5%, respectively, while those of all other subcellular compartments declined. This shift suggests an overrepresentation of nuclear proteins among the potential *N. crassa* Trx targets. Supporting this observation, 26 of the 50 most enriched proteins identified by Trx affinity chromatography are nuclear or present in both nucleus and cytoplasm (**Table 3**).

### 3.6. Kinases and transcription factors are among the most enriched proteins

Of all, the most enriched protein was the DEAD/DEAH box helicase (NCU01143), with a fold enrichment of 117 (**Table 3**). DEAD/DEAH box helicases are highly conserved in eukaryotes and play key roles in gene expression, particularly in RNA processing events such as splicing [49]. This protein belongs to the *“Gene expression and DNA/RNA-related”* category, which includes 13 other proteins among the top 50 most enriched. Notably, this group contains both PAN2 (NCU04792) and PAN3 (NCU01929), the two subunits of the deadenylation complex, shown to interact with the human TRX1 under hypoxic conditions [50]. Interestingly, this category contains also three putative transcription factors. The remaining proteins among the top 50 are distributed across five additional categories: *“Protein processing”* (8 proteins), *“Cell cycle and structure”* (4 proteins), *“Anabolism, catabolism, and energy”* (4 proteins), *“Signal transduction”* (9 proteins), and *“Uncharacterized”* (11 proteins) (**Table 3**). Strikingly, within the *“Protein processing”* category, two thiol-dependent ubiquitinyl hydrolases, NCU06294 and NCU00320, ranked second and eighth of all identified proteins, respectively, based on fold enrichment values. This category also includes four additional proteins related to the ubiquitin pathway, highlighting a potential role for redox-sensitive regulation in ubiquitin-mediated protein turnover. In the *“Signal transduction”* category, protein kinases were prominently represented, with seven out of nine proteins identified as kinases, supporting the existence of redox-sensitive signaling pathways in *N. crassa*.

To gain a comprehensive understanding of the proteins captured by Trx affinity chromatography and their potential physiological roles, we performed a Gene Ontology (GO) terms enrichment analysis among the most enriched proteins and identified the predominant terms in *“protein class”*, “*molecular function”*, and “*biological process”* categories (**Figure 2**). Regarding the *“protein class”* category, enriched terms are related to kinases, transcription factors, phospholipases, scaffold/adaptor proteins, and RNA processing factors, while unclassified proteins and transporters were underrepresented. In the *“molecular function”* category, enriched terms aligned with those found in the *“protein class”* category. For example, *“protein serine/threonine kinase activity”* for kinases, *“RNA polymerase II cis-regulatory region sequence-specific DNA binding”* for transcription factors, or *“phosphatidylinositol phospholipase C activity”* for phospholipases were overrepresented. Notably, the most enriched molecular function term was *“thioredoxin peroxidase activity”*. Additionally, three molecular function terms not captured by the protein class analysis emerged: *“chitin synthase activity”*, *“G protein activity”*, and *“iron-sulfur cluster binding”*. The *“biological process”* also highlighted ubiquitination related terms with *“Protein K11-linked ubiquitination”* being the most enriched (**Figure 2**). Altogether, eight protein types were overrepresented among the most enriched proteins: transcription factors (93 proteins), kinases (43), ubiquitination-related proteins (35), iron-sulfur cluster-related proteins (16), phospholipases (7), exonucleases (7), chitin synthases (6) and peroxiredoxins (4). We compiled a comprehensive list of all the proteins found in each of these categories (**Data Set 3**) and highlight representative ones and those mostly related fungal processes in the following paragraphs.

**Figure 2.**
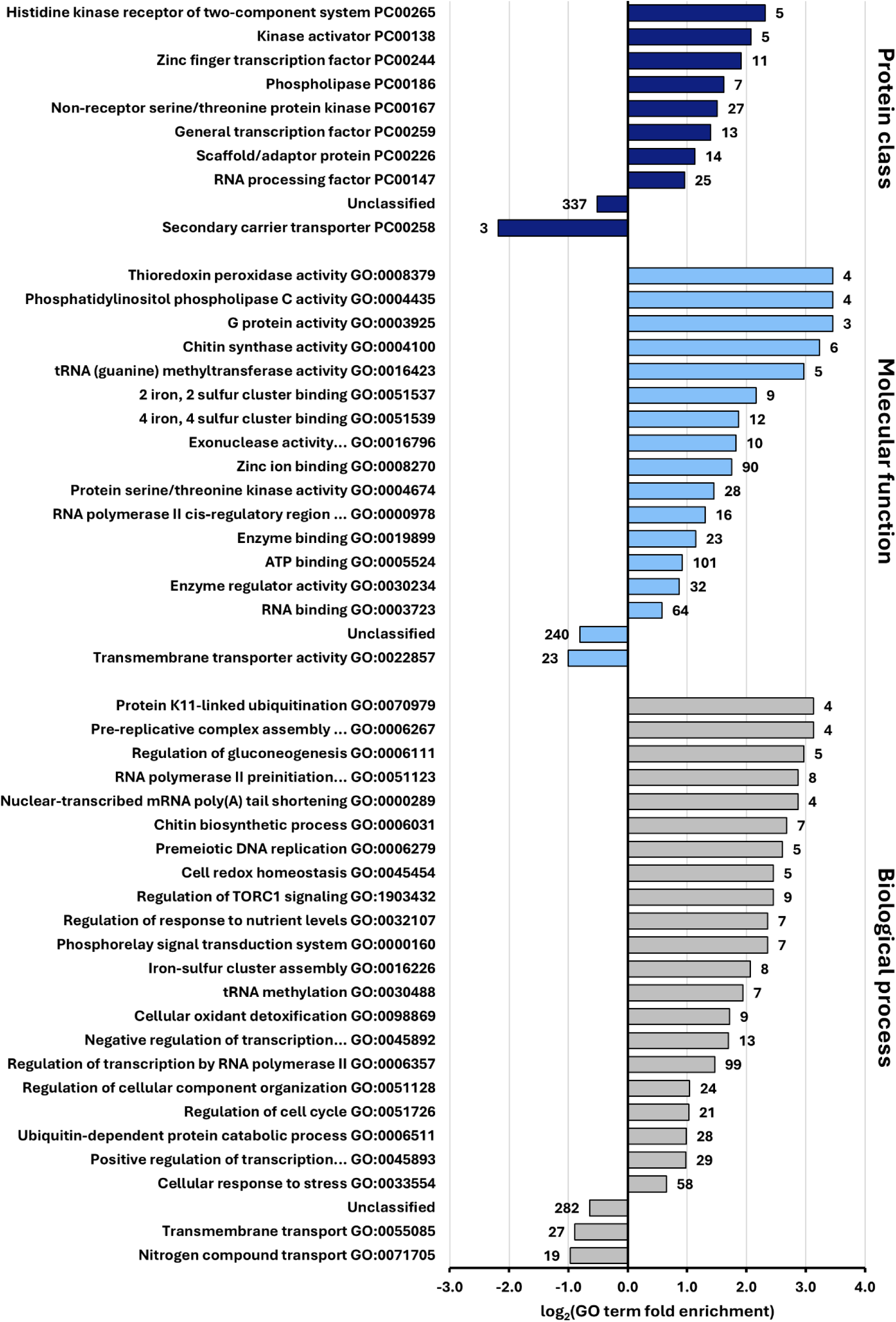
Enrichment analysis of Gene ontology terms among Trx-interacting proteins. For proteins significantly enriched during affinity chromatography (fold enrichment ≥ 4), the GO term enrichment is presented as log_2_ values for the Protein Class (*dark blue* bars), Molecular Function (*light blue* bars), and Biological Process (*gray* bars) categories. The numbers adjacent to the bars indicate the number of proteins per category. Significantly enriched GO terms were selected based on a false-discovery rate < 0.05.

Out of the five present in the genome [51], four peroxiredoxins were among the top enriched: the peroxiredoxin PRXII.1 (NCU06880) and PRXII.2 (NCU03151), the mitochondrial peroxiredoxin PRX1 (NCU06031) and the peroxiredoxin Q (NCU08931). Peroxiredoxin are ubiquitous well-known Trx targets involved in ROS scavenging and signaling [51,52], strongly supporting the relevance of fold enrichment calculations in identifying potential Trx targets using affinity chromatography.

We found a total of 93 highly enriched transcription factors (**Data Set 3**). Among them the GATA-type transcription factor SRE (NCU07728) regulates iron metabolism and is potentially activated through iron binding by conserved Cys in zinc fingers [53,54]. We also identified both the WC-1 (NCU02356) and WC-2 (NCU00902) components of the White Collar Complex (WCC), a central regulator of the circadian clock and response to light in *N. crassa* [55]. WCC has also been implicated in the regulation of CAZyme gene expression [56]. In addition, we identified two zinc-binding transcription factors involved in CAZyme regulation: CLR-1 (NCU07705), which controls the expression of cellulases [44], and PDR-1 (NCU09033), which regulates the production of pectin-degrading enzymes [57].

A total of 38 kinases and five kinase activators were identified, including 27 serine/threonine kinases and five two-component system proteins (**Data Set 3**, **Table 3**). Notably, two kinases related to conidiation were identified, the serine/threonine kinase NRC-2 (NCU01797) involved in the repression of conidia formation [58] and the MAPK kinase kinase MIK-1 (NCU02234) which plays a role in protoperithecia development [59]. We also found two other kinases participating in the regulation of protein secretion, the serine/threonine kinases MUS-59 (NCU02751) and STK-12 (NCU07378) for which gene deletions were associated with higher levels of protein secretion, among which CAZymes [60,61]. Among other remarkable kinases, the serine/threonine-protein kinase TOR (NCU05608), homolog of *S. cerevisiae* TOR2, which plays a central role in growth and development [62], and the two-component histidine kinase HCP-1 (NCU07221) were found. Interestingly in *N. crassa*, *hcp-1* was associated with the regulation of ROS production and the maintenance of cellular homeostasis in response to arthropod attack [63].

We also found 16 proteins related to iron sulfur clusters (**Data Set 3**). Among them, 12 enzymes require an iron sulfur cluster for structure or catalysis, like the three the NADH-ubiquinone oxidoreductases NUO-78 (NCU01765), NUO-51 (NCU04044) and NUO-24 (NCU01169) which are key components of the respiratory complex I [64]. The four other proteins are potentially involved in iron-sulfur cluster assembly, such as NAR-1 (NCU03204) whose homolog in *S. cerevisiae* was shown to transfer an Fe/S cluster to apoproteins thanks to conserved Cys residues [65].

Interestingly, out of the seven chitin synthases encoded in *N. crassa* genome [66], we found six among the top-enriched proteins (**Data Set 3**). These enzymes are essential for cell wall biosynthesis, morphogenesis, and development of the fungus, and our results suggest potential redox regulation.

Altogether, the identity of the proteins with high fold enrichment values (≥ 4) reveal a diverse set of redox-sensitive candidate Trx targets in *N. crassa*. In particular, the high number of kinases and transcription factors supports the existence of redox-regulated signaling pathways in the fungus.

### 3.7. Multiprotein complexes are enriched by Trx affinity chromatography

We noticed that numerous candidate Trx targets were annotated as subunits or components of multiprotein complexes. To obtain a broader view of potential complex-associated proteins within our dataset, we performed Gene Ontology (GO) enrichment analysis for the “Cellular Component” category. This analysis highlighted nine multiprotein complexes, each represented by three or more components (**Figure 3**). All the components were isolated for five complexes: the three subunits of the GATOR1 complex, a negative regulator of mTORC1 signaling that promotes autophagy under amino acid starvation [67], the four subunits of the Ndc80 complex, required for accurate chromosome segregation during mitosis [68], the three subunits of the L-cysteine desulfurase complex, which mobilizes sulfur from Cys to enable biosynthesis of iron-sulfur clusters and sulfur-containing cofactors [69], the four components of the GID complex, a conserved E3 ubiquitin ligase involved in targeted degradation of gluconeogenic enzymes [70] and the three subunits of the protein phosphatase 4 complex, which plays roles in DNA repair, cell cycle control, metabolism, and immune responses [71]. We also found ten subunits of the transcription factor TFIID complex, a central regulator of RNA polymerase II transcription [72], eight subunits of the mRNA cleavage and polyadenylation specificity factor complex, critical for mRNA maturation [73], eight components of the histone deacetylase complex, involved in chromatin condensation and gene silencing [74] and four subunits of the Smc5/6 complex, which contributes to genome stability during DNA replication and repair [75]. We cannot determine whether these proteins were isolated individually, or co-purified through interactions with redox-sensitive subunits of complexes. However, we observed that eight protein components lacked Cys, suggesting they were co-purified through interactions with redox-sensitive subunits. This highlights that Trx affinity chromatography could catch multiprotein complexes in addition to individual redox-regulated Trx targets.

**Figure 3.**
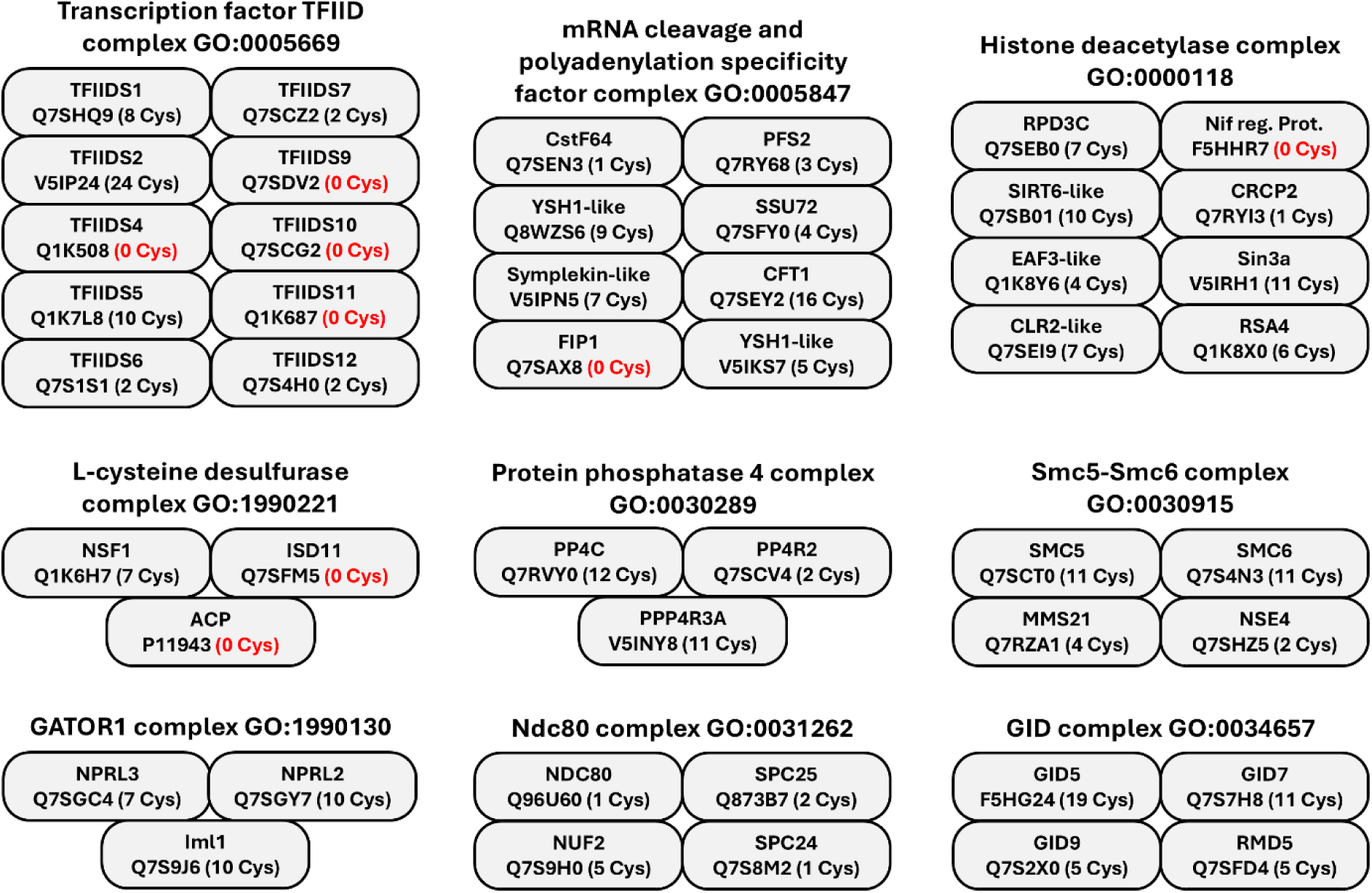
Components of multiprotein complexes isolated by Trx affinity chromatography. Components of multiprotein complexes identified in the Gene Ontology (GO) category “Cellular Components”, among the proteins with fold enrichment ≥ 4. Significantly enriched GO terms were selected based on a false-discovery rate < 0.05. The number of Cys is indicated.

## 4. Discussion

In this study, we identified the components of the Trx system in *N. crassa* and used affinity chromatography to uncover potential redox-regulated Trx target proteins. The Trx system of *N. crassa* displays similarities with the one of *S. cerevisae*, with three Trx and a TR. This low number of genes is consistent with observations in other fungal species [22,51]. Similarly, fungi also possess relatively few genes encoding other types of thiol-oxidoreductases, such as peroxiredoxins and methionine sulfoxide reductases [51,76], suggesting that, as in animals and bacteria, the overall number of thiol-oxidoreductase genes is generally low in fungi. However, our analysis revealed discrepancies in predicted subcellular localizations. TRX1, TRX2 and TRX3 are predicted to localize to the endoplasmic reticulum (ER), cytoplasm, and mitochondria, respectively, whereas TR is predicted to reside in the cytoplasm or the peroxisome (**Table 1**). If these predictions are accurate, they would prevent the physical interaction between TR and TRX1 or TRX3, which is essential for maintaining their reduced state. Notably, the predicted localization of TRX1 to the ER is unexpected, as, to our knowledge, no Trx was reported in the ER, and its overexpression in this compartment induced cytotoxicity in human cells [77]. It will be necessary to precisely determine the subcellular localization of these proteins, taking into consideration that it can be tissue-dependent and modified in response to environmental cues [78,79]. Additionally, TRX1 has a C-terminal extension conserved in numerous fungi, but only in two classes of *Pezizomycotina* (*Leotiomycetes* and *Sordariomycetes*) (**Table S1**). As *N. crassa* also belongs to the *Sordariomycetes*, the confined occurrence of the C-terminal extension in these phyla suggests a specialized function potentially linked to lineage-specific physiological roles.

Applying Trx affinity chromatography to *N. crassa*, we identified 1,998 proteins, corresponding to approximately 19% of the total proteome [24]. Similarly high number of Trx interacting proteins was previously reported for the algae *Chlamydomonas reinhardtii* [6]. While early studies identified restricted numbers of potential Trx targets [80], the experiments made with the algae and ours identified high numbers of proteins very likely because of the major advancements made in mass spectrometry sensitivity. As many large-scale approaches, Trx affinity chromatography can yield false positives and may fail to capture some genuine Trx targets [80]. Based on the nature of their interaction with Trx, the proteins identified can be categorized into four groups. First, the genuine physiological Trx targets, i.e. the proteins that are redox-regulated and form transient intermolecular disulfide bridges with a Trx. Second, the proteins that interact with the Trx, but not through intermolecular disulfide. They can be regulators or partners but are not necessarily redox-regulated proteins. Third, redox-sensitive proteins that are not targeted by Trx, but potentially regulated by alternative systems such as glutaredoxins. Fouth, non-specific interactors that are false positives recovered due to high abundance or unspecific non-covalent associations, but also potentially proteins that associate with a Trx target as part of larger multiprotein complexes. Additionally, genuine Trx targets may be missed due to low abundance or because they were not oxidized under the experimental conditions. Because of these flaws in the affinity chromatography approach and the high number of proteins identified, we applied a fold enrichment calculation to rank the proteins in the eluates. The comparison of the proteins identified in each eluate ranked by fold enrichment gave very similar results, with approximately one-third of the proteins with fold enrichment below one, one-third with value between 1 and the mean 4, and the last third with value above 4 and up to 117 for the most enriched (**Tables 2, 3**). The fold enrichment value of proteins with at least one Cys was significantly higher than the one of proteins without Cys, validating this approach. It should be noted that some proteins without Cys were highly enriched because of their presence in multiprotein complexes (**Figure 3**). To our knowledge, such quantitative analysis and rankings were never applied for Trx affinity chromatography. Proteins with fold enrichment values below one are unlikely to be Trx targets, although we cannot exclude the possibility that targets were not oxidized during the assay. In contrast, proteins with fold enrichment values above one were more abundant after affinity chromatography than in the reference proteome and likely include genuine Trx targets in *N. crassa*. Focusing on the proteins with fold enrichment values equal or above four showed strong redundancy between the sets of proteins retrieved with each Trx, with most proteins found in all eluates. Such redundancy and lack of specificity for Trx were previously reported [6,16,81]. This suggests that *N. crassa*’s Trx could potentially reduce the same targets *in vitro* and that the specificity might arise from the co-localization of the Trx and its target within the same subcellular compartment.

Among the most enriched proteins, we found homologs of known Trx targets in other organisms and uncovered a wealth of potential fungal Trx targets or redox regulated proteins. Interestingly, nuclear proteins were overrepresented among them, with notably almost a hundred transcription factors (**Data Set 3**). Once again, this raises a potential inconsistency between the subcellular localization of the Trx and their putative targets, as transcription factors are active in the nucleus. However, as previously shown for Yap1, that we found in our experiment, redox-regulated transcription factor can shuttle from the cytoplasm to the nucleus [82]. Moreover, we found more than 40 kinases, suggesting that numerous signaling pathways might be redox regulated by Trx in *N. crassa*. Interestingly, we identified three transcription factors (CLR-1, PDR-1, and WCC) and two kinases (MUS-59 and STK-12) known to regulate CAZyme production and protein secretion [44,55,57,60,61]. This finding suggests that key processes underpinning the saprotrophic lifestyle may be subject to redox regulation. Supporting this idea, hydrogen peroxide was shown to modulate WCC activity, likely through redox-sensitive PAS or LOV domains [83].

Future research will aim to precisely determine the subcellular localization of the Trx in *N. crassa*, as our data suggests potential discrepancies between Trx and TR localization that could affect redox regulation. Additionally, experimental validation of redox-sensitive interactions will be essential to distinguish genuine Trx targets from indirect or non-specific interactors identified through affinity chromatography. Particular attention will be given to the nuclear and signaling proteins uncovered in our study, including numerous transcription factors and kinases, which suggest a broad role for Trx in the regulation of gene expression and signaling pathways. Ultimately, elucidating the redox-regulatory network mediated by Trx in *N. crassa* will provide valuable insights into fungal biology, particularly in the context of CAZymes production and plant cell wall degradation.

## Supporting information

Data Set 1

Data Set 2

Data Set 3

## CRediT authorship contribution statement

**Lucia Bidondo:** Conceptualization, Methodology, Validation, Formal analysis, Investigation, Data Curation, Writing - Original Draft, Writing - Review & Editing, Visualization. **Elodie Drula:** Investigation, Data Curation. **Jean-Charles Gaillard:** Formal analysis, Investigation. **Jean Armengaud:** Formal analysis, Resources, Data Curation, Writing - Review & Editing. **Jean-Guy Berrin:** Resources, Writing - Review & Editing, Project administration, Funding acquisition. **Marie-Noëlle Rosso:** Conceptualization, Methodology, Validation, Investigation, Resources, Writing - Original Draft, Writing - Review & Editing, Supervision, Funding acquisition. **Lionel Tarrago:** Conceptualization, Methodology, Validation, Formal analysis, Investigation, Resources, Data Curation, Writing - Original Draft, Writing - Review & Editing, Visualization, Supervision, Funding acquisition.

## Data availability

The mass spectrometry proteomics data have been deposited to the ProteomeXchange Consortium via the PRIDE partner repository with the dataset identifier PXD063284 and 10.6019/PXD063284.

## Funding

This study received funding from the Novo Nordisk Foundation grant NNF20OC0059697 - OxyMiST project. This work received support from the French government under the France 2030 investment plan, as part of the Initiative d’Excellence d’Aix-Marseille Université – A*MIDEX (AMX-21-PEP-028) for L.T.

## Acknowledgments

The authors kindly thank the support team of Aix Marseille Univ., INRAE, Biodiversité et Biotechnologies Fongiques – Chantal Parodi-Negri, Naura Thibeau and Christophe Boyer, and all the members of the OxyMiST project for fruitful discussions.

## Notes

### Competing Interest Statement

The authors have declared no competing interest.

