## Supporting information for "Global mapping of thioredoxin interacting proteins in *Neurospora crassa*"

**From**

[1]INRAE, Aix Marseille Univ., BBF, Biodiversité et Biotechnologie Fongiques, Marseille, France.

[2]CNRS, Aix Marseille Univ, AFMB, USC1408, Marseille, France

[3]Université Paris-Saclay, CEA, INRAE, Département Médicaments et Technologies pour la Santé (DMTS), SPI, 30200 Bagnols-sur-Cèze, France.

### SUPPORTING INFORMATION

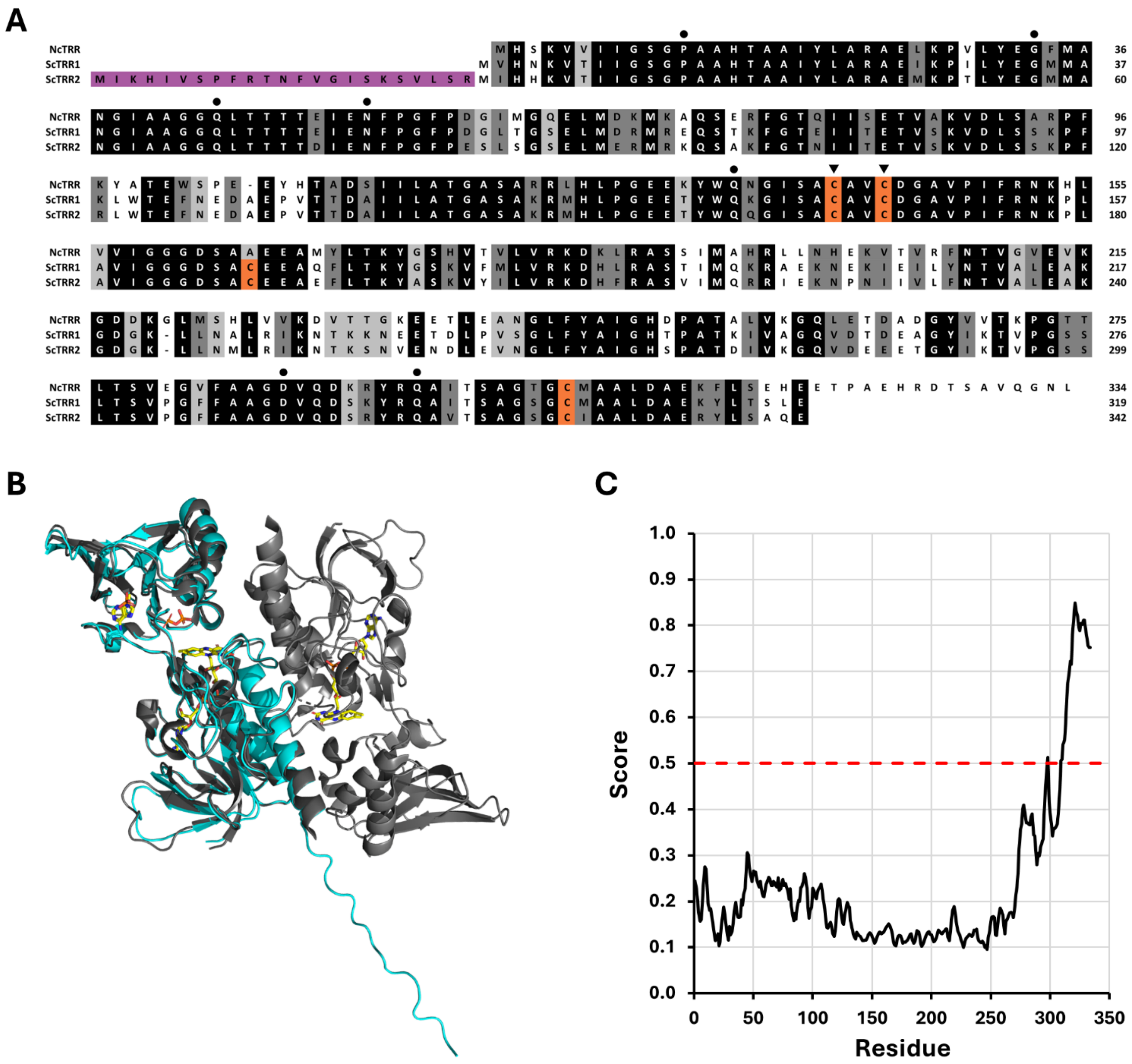

**Figure S1. Protein sequence and 3D model of *N. crassa* TR aligned with *S. cerevisiae* homologs.**

**A)** Protein sequence alignment of *N. crassa* TRR (NcTRR; UniProt#P51978) with *S. cerevisisae* TRR1 (ScTRR1; Uniprot# P29509) and TRR2 (ScTRR2; Uniprot# P38816). NcTRR shares 65.8% and 59.6% identity, and 78.6% and 71.6% similarity with ScTRR1 and ScTRR2, respectively. Percentages were calculated using pairwise alignment using Emboss Needle with default parameters (https://www.ebi.ac.uk/jdispatcher/psa/emboss_needle).

**B)** Structural alignment of the predicted 3D model (AlphaFold 2.0) of NcTRR (in *cyan*) with the crystallographic structure of ScTRR1 (PDB# 3D8X), in *gray*, performed using Open-source PyMol 3.1. NAD and FAD cofactors are shown in sticks.

**C)** Prediction of disordered regions in the NcTRR primary sequence using IUPred (https://aiupred.elte.hu/). Each residue was assigned a disorder probability score from 0 to 1, with values above 0.5 (*dashed red* line) indicating a high likelihood of structural disorder.

A

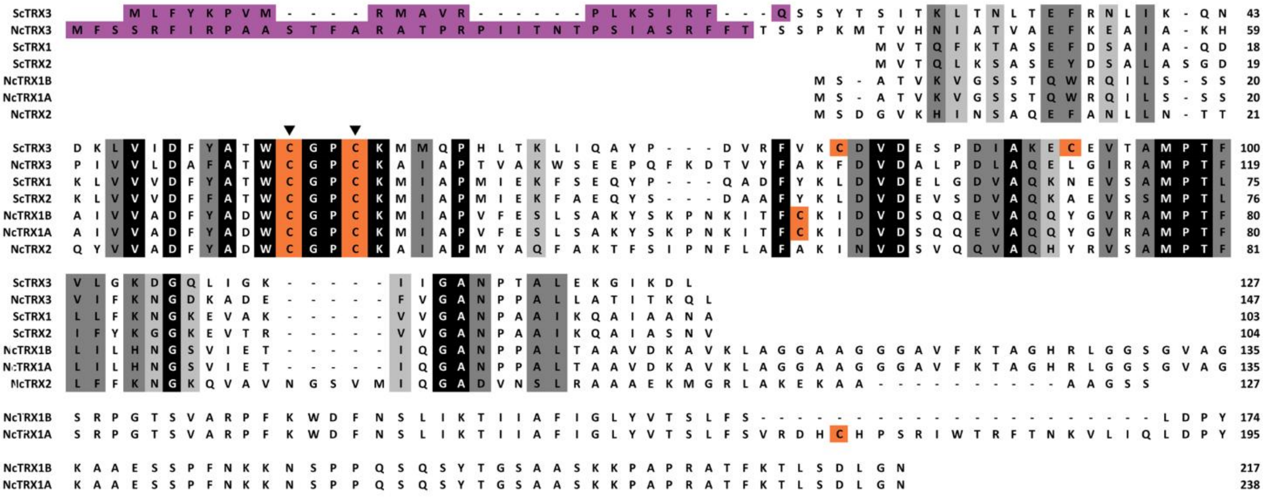

B

| | NcTRX1B | NcTRX2 | NcTRX3 | ScTRX1 | ScTRX2 | ScTRX3 |
|---|---|---|---|---|---|---|
| NcTRX1A | 91.2 (91.2) | 23.9 (32.0) | 15.5 (23.1) | 18.1 (26.9) | 17.2 (25.9) | 16.1 (24.9) |
| NcTRX1B | | 26.1 (35.0) | 16.8 (25.0) | 19.8 (29.5) | 18.8 (28.4) | 17.5 (27.1) |
| NcTRX2 | | | 24.6 (37.1) | 34.4 (46.6) | 35.7 (51.9) | 22.8 (39.6) |
| NcTRX3 | | | | 33.8 (44.6) | 28.4 (43.9) | 36.7 (51.7) |

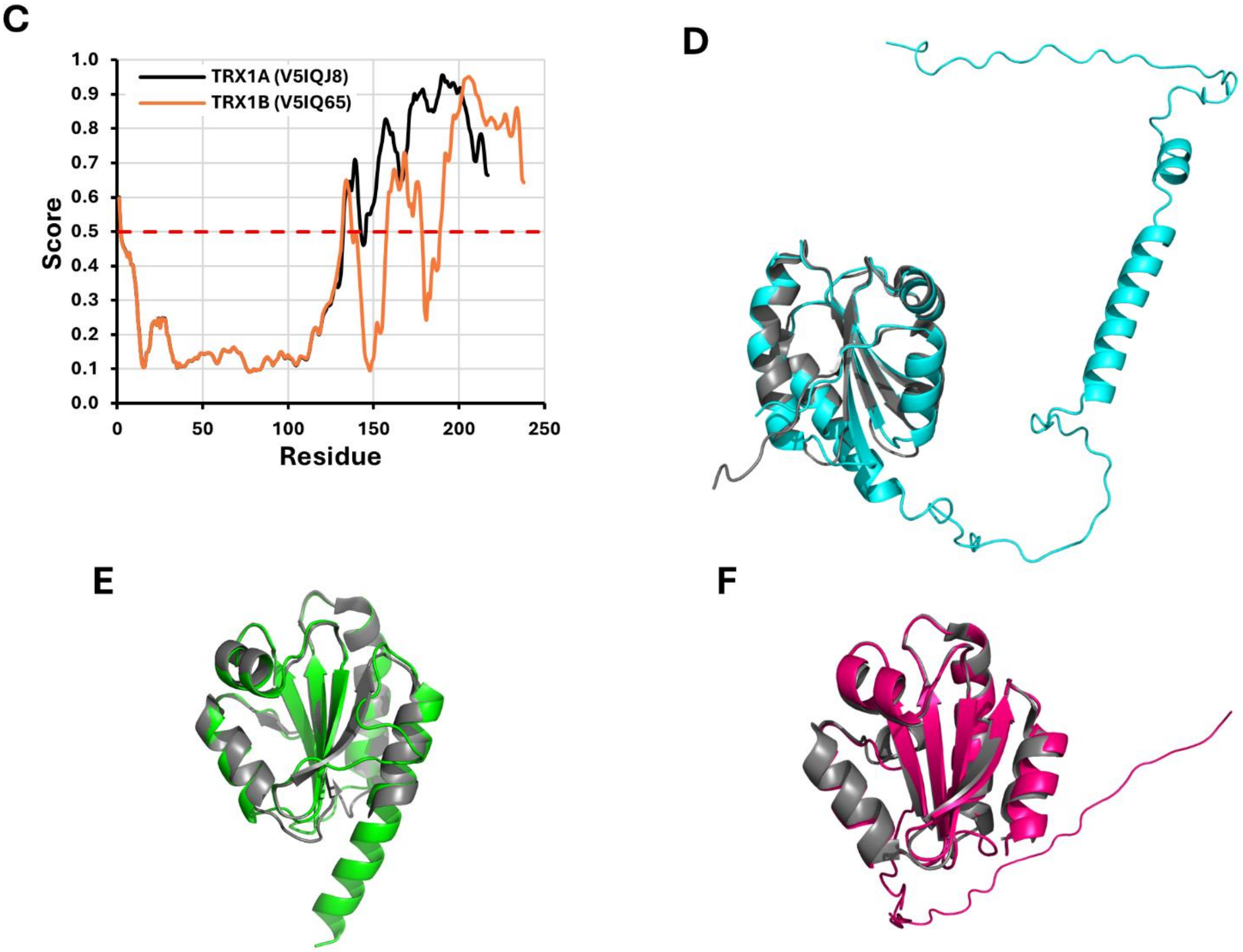

**Figure S2. Protein sequence and 3D model of *N. crassa* Trx aligned with *S. cerevisiae* homologs.**

**A)** Protein sequence alignment of *N. crassa* TRX1A (NcTRX1A; UniProt# V5IQ65), *N. crassa* TRX1B (NcTRX1B; Uniprot# V5IQJ8), *N. crassa* TRX2 (NcTRX2; Uniprot# P42115) and *N. crassa* TRX3 (NcTRX3; Uniprot# F5HBN4) with *S. cerevisiae* TRX1 (ScTRX1; Uniprot# P22217), TRX2 (ScTRX2; Uniprot# P22803) and TRX3 (ScTRX3; Uniprot# P25372).

**B)** Pairwise sequence identity and similarity percentages between NcTrx and ScTrx proteins, calculated using EMBOSS Needle with default parameters (https://www.ebi.ac.uk/jdispatcher/psa/emboss_needle).

**C)** Prediction of intrinsically disordered regions in NcTRX1A and NcTRX1B primary sequences using IUPred (https://aiupred.elte.hu/). Each residue is assigned a disorder probability score from 0 to 1, with scores above 0.5 (indicated by the *dashed red* line) denoting predicted disorder.

**D-F)** Structural alignment of *N. crassa* Trx 3D models (AlphaFold 2.0) with the experimentally determined *S. cerevisiae* Trx structures:: **C)** TRX1 (NcTRX1 in *cyan*, ScTRX1 PDB# 3F3Q in *gray*), **D)** TRX2 (NcTRX2 in *green*, ScTRX2 PDB# 2FA4 in *gray*), **E)** TRX3 (NcTRX3 in *pink*, ScTRX3 PDB# 2OE3 in *gray*).

**A**

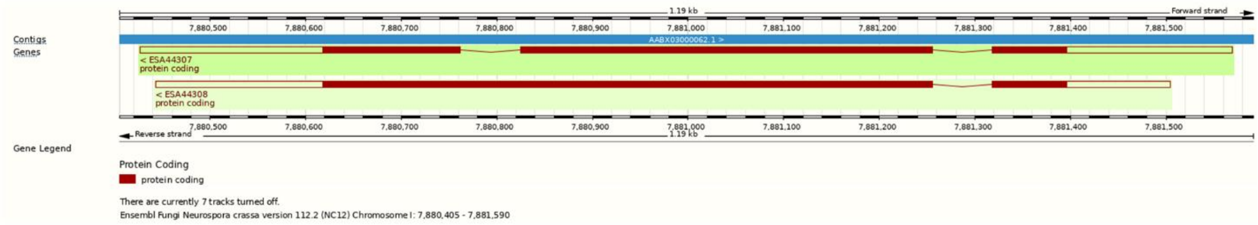

**B**

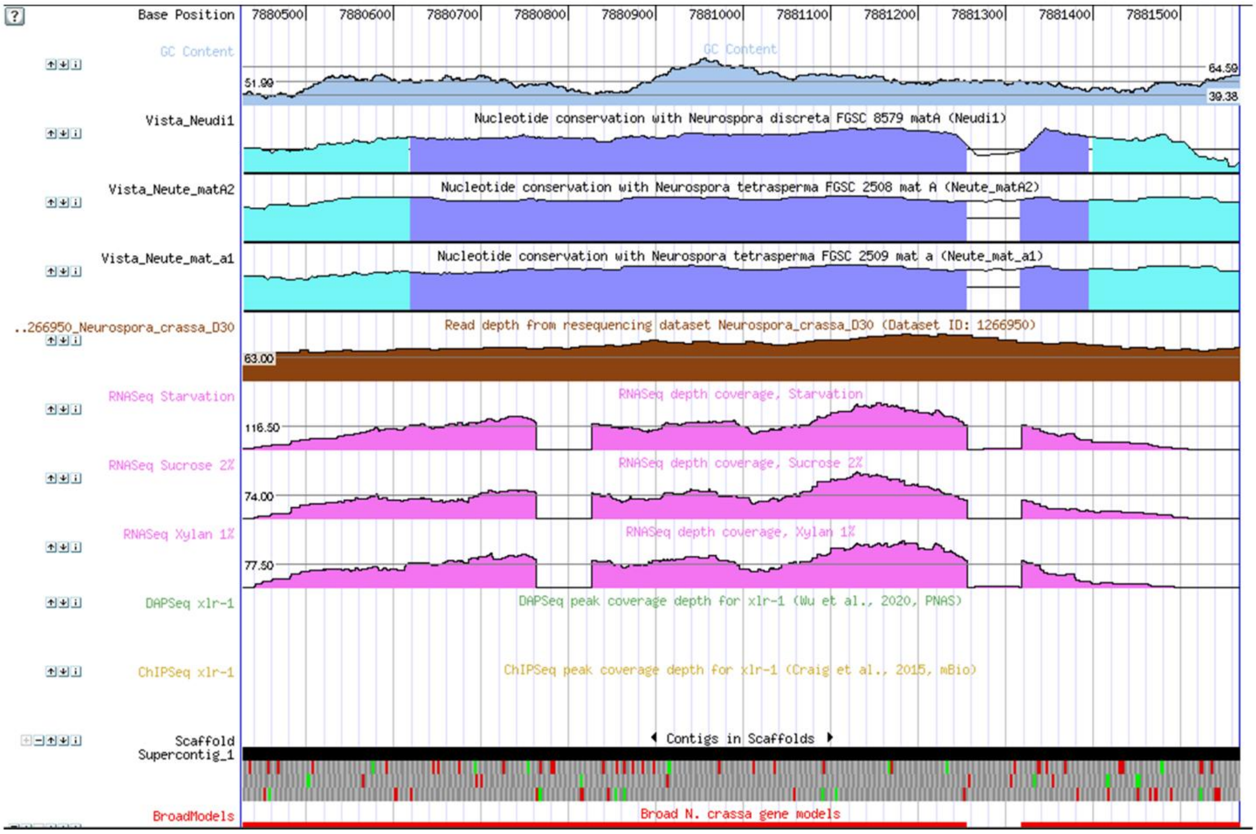

**Figure S3. Structure of the NCU00598 gene coding for TRX1 and RNAseq coverage.**

**A)** Screenshot of the Ensembl Fungi website for the region of NCU00598 gene showing the alternative splicing leading to the two TRX1 variants (http://fungi.ensembl.org/Neurospora_crassa/Location/View?db=core;g=NCU00598;r=I:7880405-7881590;t=ESA44307).

**B)** Screenshot of the genome browser of the MycoCosm website corresponding to the region of NCU00598 gene showing the coverage of RNAseq data (in *pink*) grown under starvation, on 2% sucrose or on 1% xylan (https://mycocosm.jgi.doe.gov/cgi-bin/browserLoad/?db=Neucr2&position=Supercontig_1:7880427-7881568).

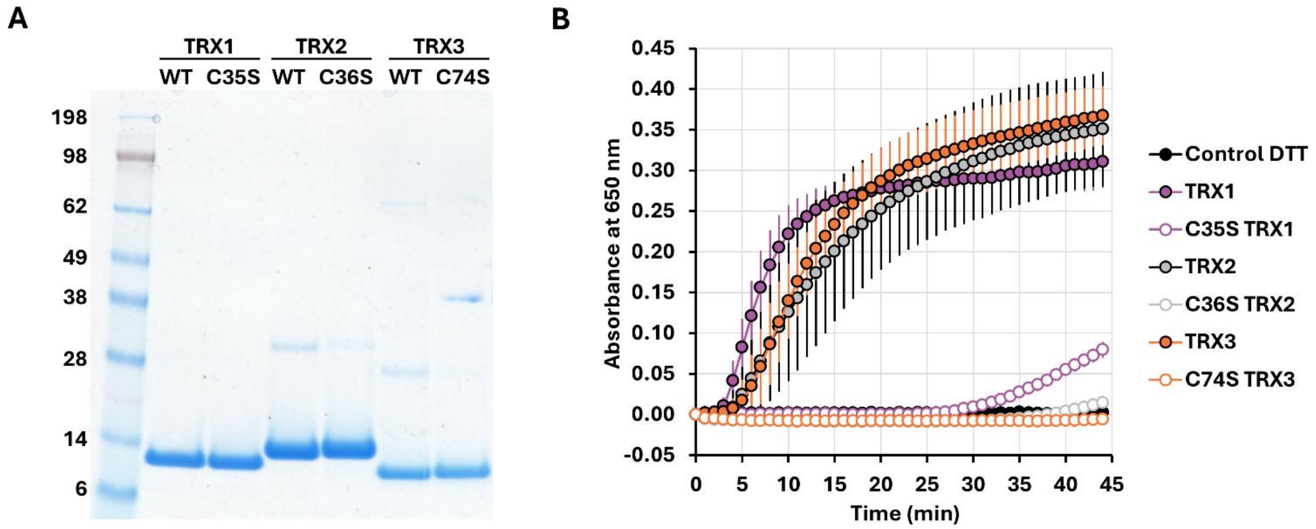

**Figure S4. Purification of wild-type and Cys-to-Ser mutants of Trx and insulin reduction assay.**

**A)** After production and purification as described in material and methods, 5 µg of wild-type and Cys-to-Ser mutant of TRX1, TRX2 and TRX3 were analyzed by SDS-PAGE.

**B)** Trx activity was determined following insulin precipitation at 650 nm with 5 µM Trx and 1 mM DTT in 0.1 M $K_2HPO_4$, 2 mM EDTA and pH 7.5.

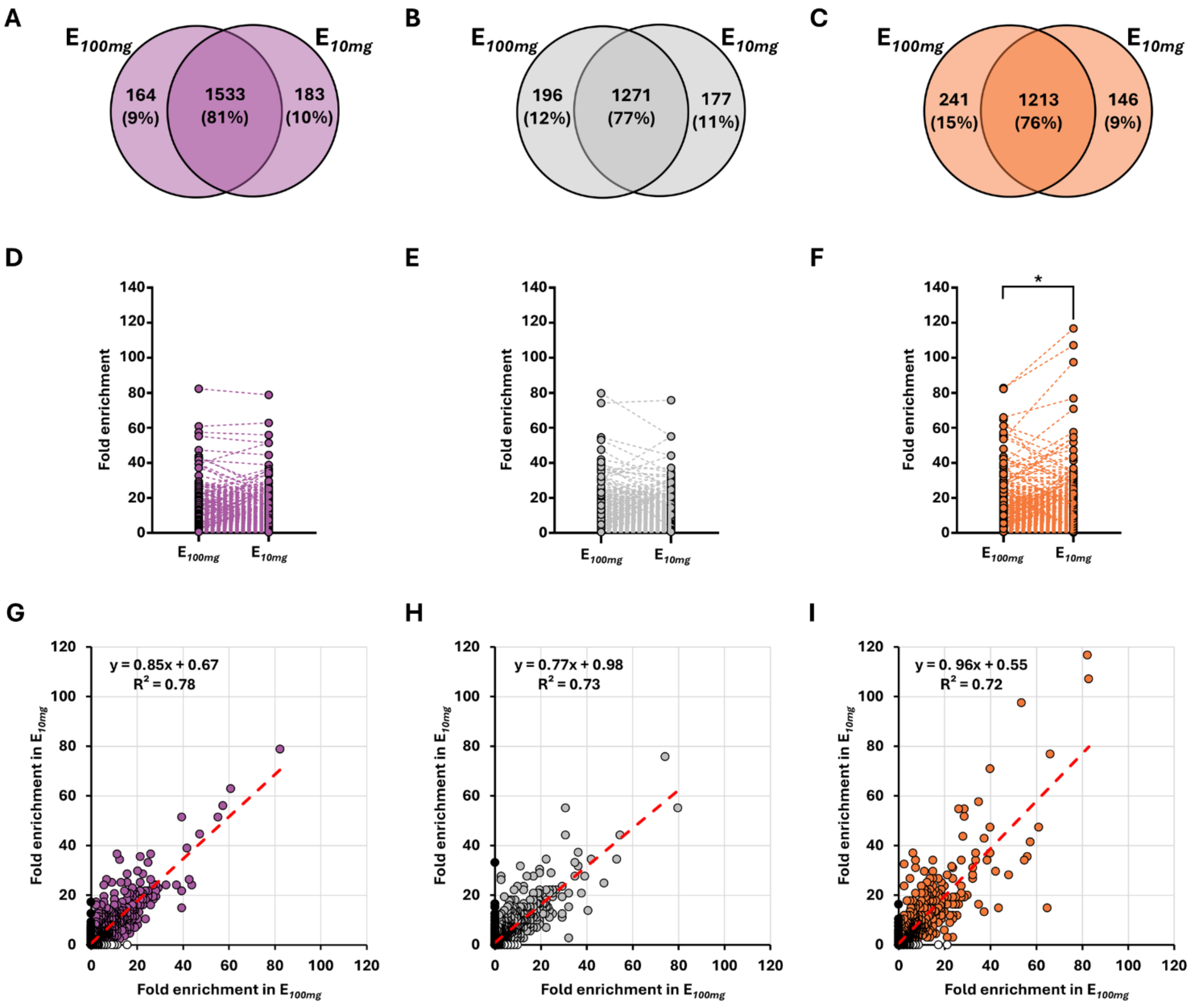

**Figure S5. Comparison of fold enrichment in $E_{100mg}$ vs $E_{10mg}$ eluates for each Trx.**

**A, B, C)** Venn diagram of proteins found in both $E_{100mg}$ and $E_{10mg}$ for TRX1, TRX2 and TRX3, respectively.

**D, E, F)** Bar graph of fold enrichment for proteins found in both $E_{100mg}$ and $E_{10mg}$ after TRX1, TRX2 and TRX3 affinity chromatography, respectively. Statistical analysis was performed using a Wilcoxon signed rank test (**P* = 0.0173).

**G, H, I)** For all proteins, the fold enrichment in $E_{10mg}$ was plotted as a function of the fold enrichment in $E_{100mg}$ for TRX1, TRX2 and TRX3, respectively. Linear regression is presented in *red*, with equation and $R^2$ values indicated. Each colored dot (*purple*, *gray* or *orange*) represents a protein found in both $E_{100mg}$ and $E_{10mg}$, while *white* and *black* dots represent protein found only in $E_{100mg}$ or $E_{10mg}$, respectively.

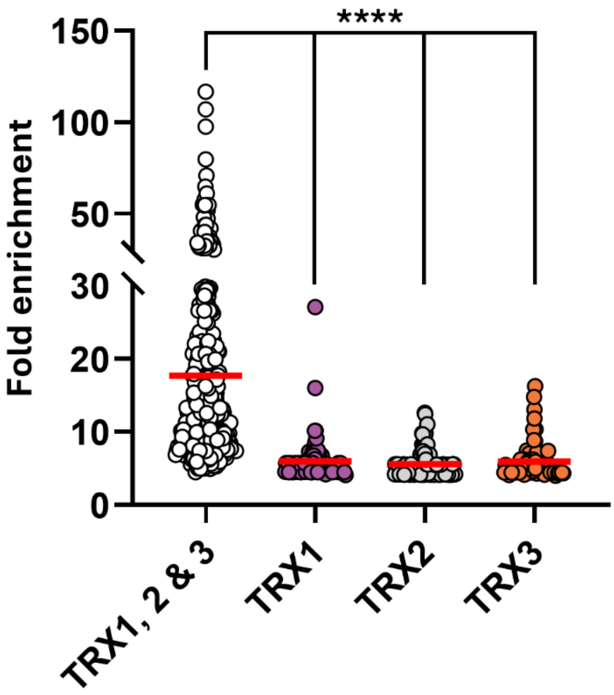

**Figure S6. Fold enrichment of proteins found common among all eluted proteins (*"TRX1, 2 & 3"*) and those unique to TRX1, TRX2, or TRX3 eluates (*"TRX1"*, *"TRX2"* and *"TRX3"*, respectively).**

Each dot represents a protein, with the mean fold enrichment indicated by a horizontal *red* bar. Only proteins with a fold enrichment ≥ 4 were considered (Data Set 1). These groups correspond to those of each group in Figure 1C, D. The number of proteins were 386, 83, 142 and 81 for TRX1, 2 & 3, TRX1, TRX2 and TRX3, respectively. The mean fold enrichment was ~18 for TRX1, 2 & 3 and ~6 for the three other groups. Statistical analysis was performed using Mann-Whitney test (****$P < 0.0001$).

**Table S1. Fungal thioredoxins with a C-terminal extension similar to that of *N. crassa* TRX1.**

| Class | Order | Family | Genus | Species | Uniprot Accession |
|---|---|---|---|---|---|
| Leotiomycetes | Erysiphales | Erysiphaceae | Blumeria | *Blumeria graminis* f. sp. hordei (strain DH14) | N1JHN9 |
| | | | | *Blumeria graminis* f. sp. tritici | A0A9X9MGA8 |
| | | | | *Blumeria graminis* f. sp. tritici 96224 | A0A061HMD0 |
| | | | Erysiphe | *Erysiphe neolycopersici* | A0A420HZ12 |
| | | | | *Erysiphe pulchra* | A0A2S4PW70 |
| | | | Golovinomyces | *Golovinomyces cichoracearum* | A0A420J8M4 |
| | Helotiales | Amorphothecaceae | Amorphotheca | *Amorphotheca resinae* ATCC 22711 | A0A2T3AV02 |
| | | Dermateaceae | Coleophoma | *Coleophoma cylindrospora* | A0A3D8RUC3 |
| | | | Marssonina | *Marssonina coronariae* | A0A218YW76 |
| | | Drepanopezizaceae | Drepanopeziza | *Marssonina brunnea* f. sp. multigermtubi (strain MB_m1) | K1X8Z7 |
| | | Helotiaceae | Glarea | *Glarea lozoyensis* (strain ATCC 20868 / MF5171) | S3D9I4 |
| | | | Hymenoscyphus | *Hymenoscyphus albidus* | A0A9N9LGQ1 |
| | | | | *Hymenoscyphus fraxineus* | A0A9N9L3L2 |
| | | Helotiales *incertae sedis* (no rank) | Amylocarpus | *Amylocarpus encephaloides* | A0A9P7YB35 |
| | | | Cadophora | *Cadophora* sp. M221 | A0A8H7XHX6 |
| | | | | *Cadophora malorum* | A0A8H7WGY0 |
| | | | Rhexocercosporidium | *Rhexocercosporidium* sp. MPI-PUGE-AT-0058 | A0A9P9S0J3 |
| | | Hyaloscyphaceae | Hyaloscypha | *Hyaloscypha bicolor* E | A0A2J6T3M4 |
| | | | | *Hyaloscypha variabilis* (strain UAMH 11265 / GT02V1 / F) | A0A2J6RLE0 |
| | | | | *Hyaloscypha hepaticicola* | A0A2J6PDS6 |
| | | Hyphodiscaceae | Hyphodiscus | *Hyphodiscus hymeniophilus* | A0A9P7AYY4 |
| | | Lachnaceae | Lachnellula | *Lachnellula suecica* | A0A8T9CQS7 |
| | | | | *Lachnellula arida* | A0A8T9BHT6 |
| | | | | *Lachnellula occidentalis* | A0A8H8UBU9 |
| | | | | *Lachnellula subtilissima* | A0A8H8U6Q1 |
| | | | | *Lachnellula hyalina* | A0A8H8R0N2 |
| | | | | *Lachnellula cervina* | A0A7D8YVE0 |
| | | | | *Lachnellula willkommii* | A0A559M161 |
| | | Mollisiaceae | Mollisia | *Mollisia scopiformis* | A0A194X1G3 |
| | | | Phialocephala | *Phialocephala subalpina* | A0A1L7X0Z7 |
| | | Pezizellaceae | Calycina | *Calycina marina* | A0A9P7Z198 |
| | | Pleuroascaceae | Venustampulla | *Venustampulla echinocandica* | A0A370TKU9 |
| | | Ploettnerulaceae | Rhynchosporium | *Rhynchosporium commune* | A0A1E1KUP3 |
| | | Rutstroemiaceae | Rutstroemia | *Rutstroemia* sp. NJR-2017a WRK4 | A0A2S7R1C4 |
| | | | | *Rutstroemia* sp. NJR-2017a BVV2 | A0A2S7QCX2 |
| | | | | *Rutstroemia* sp. NJR-2017a BBW | A0A2S7P890 |

**Table S1.** ***(continued)***

| Class | Order | Family | Genus | Species | Uniprot Accession |
|---|---|---|---|---|---|
| Leotiomycetes | Helotiales | Sclerotiniaceae | Botryotinia | *Botryotinia convoluta* | A0A4Z1IU82 |
| | | | | *Botryotinia narcissicola* | A0A4Z1HAB8 |
| | | | | *Botryotinia calthae* | A0A4Y8D6X3 |
| | | | Botrytis | *Botryotinia fuckeliana* (strain T4) (*Botrytis cinerea*) | G2XZF2 |
| | | | Botrytis | *Botrytis fragariae* | A0A8H6B3C3 |
| | | | Botrytis | *Botrytis elliptica* | A0A4Z1K2F6 |
| | | | Botrytis | *Botryotinia fuckeliana* (strain B05.10) (*Botrytis cinerea*) | A0A384J690 |
| | | | Monilinia | *Monilinia vaccinii-corymbosi* | A0A8A3P120 |
| | | | | *Monilinia laxa* | A0A5N6KF75 |
| | | | | *Monilinia fructicola* | A0A5M9K8H1 |
| | | | Sclerotinia | *Sclerotinia borealis* (strain F-4128) | W9CTM8 |
| | | | | *Sclerotinia nivalis* | A0A9X0DRB6 |
| | | | | *Sclerotinia trifoliorum* | A0A8H2W0D1 |
| | | Tricladiaceae | Cudoniella | *Cudoniella acicularis* | A0A8H4RQU3 |
| | | unclassified Helotiales (no rank) | | *Helotiales* sp. DMI_Dod_QoI | A0A8S9BPJ6 |
| | Leotiomycetes *incertae sedis* (no rank) | Myxotrichaceae | Oidiodendron | *Oidiodendron maius* (strain Zn) | A0A0C3D8M0 |
| | | Pseudeurotiaceae | Pseudogymnoascus | *Pseudogymnoascus destructans* (strain ATCC MYA-4855 / 20631-21) | L8FQG1 |
| | | | | *Pseudogymnoascus verrucosus* | A0A1B8GJF6 |
| | | | | *Pseudogymnoascus sp. 05NY08* | A0A1B8F6W5 |
| | | | | *Pseudogymnoascus* sp. 23342-1-I1 | A0A1B8DXP6 |
| | | | | *Pseudogymnoascus* sp. 24MN13 | A0A1B8CY58 |
| | | | | *Pseudogymnoascus* sp. WSF 3629 | A0A1B8C7E4 |
| | | | | *Pseudogymnoascus* sp. VKM F-4519 (FW-2642) | A0A094IWZ0 |
| | | Pseudeurotiaceae | Pseudogymnoascus | *Pseudogymnoascus* sp. VKM F-4515 (FW-2607) | A0A094EKJ6 |
| | | | | *Pseudogymnoascus* sp. VKM F-4516 (FW-969) | A0A094D7C2 |
| | | | | *Pseudogymnoascus* sp. VKM F-4281 (FW-2241) | A0A094D6T6 |
| | | | | *Pseudogymnoascus* sp. VKM F-4513 (FW-928) | A0A094BYV8 |
| | | | | *Pseudogymnoascus* sp. VKM F-4246 | A0A093Z157 |
| | | | | *Pseudogymnoascus* sp. VKM F-3808 | A0A093YGM5 |
| | | | | *Pseudogymnoascus* sp. VKM F-3557 | A0A093XR72 |
| | unclassified Leotiomycetes (no rank) | | | *Leotiomycetes* sp. MPI-SDFR-AT-0126 | A0A9P9P9Z9 |

**Table S1.** ***(continued)***

| Class | Order | Family | Genus | Species | Uniprot Accession |
|---|---|---|---|---|---|
| Sordariomycetes | Coniochaetales | Coniochaetaceae | Coniochaeta | *Coniochaeta* sp. 2T2.1 | A0A5N5MYG3 |
| | Diaporthales | Cryphonectriaceae | Cryphonectria-Endothia species complex (no rank) | *Cryphonectria parasitica* EP155 | A0A9P5CSY6 |
| | | Diaporthaceae | Diaporthe | *Diaporthe helianthi* | A0A2P5HYI2 |
| | | | Diaporthe | *Diaporthe ampelina* | A0A0G2HW45 |
| | | Gnomoniaceae | Gnomoniopsis | *Gnomoniopsis smithogilvyi* | A0A9W9CTP3 |
| | | Schizoparmaceae | Coniella | *Coniella lustricola* | A0A2T3A9H4 |
| | | Valsaceae | Cytospora | *Cytospora leucostoma* | A0A423XJN7 |
| | | | Valsa | *Valsa malicola* | A0A423X3U5 |
| | | | | *Valsa sordida* | A0A423VD77 |
| | | | | *Valsa mali* var. pyri (nom. inval.) | A0A194V4Y1 |
| | Glomerellales | Glomerellaceae | Colletotrichum | *Colletotrichum gloeosporioides* (strain Cg-14) | T0L2G4 |
| | | | | *Colletotrichum orbiculare* (strain 104-T / ATCC 96160 / CBS 514.97 / LARS 414 / MAFF 240422) | N4VX39 |
| | | | | *Colletotrichum fructicola* (strain Nara gc5) | L2FS47 |
| | | | | *Colletotrichum graminicola* (strain M1.001 / M2 / FGSC 10212) | E3QP06 |
| | | | | *Colletotrichum noveboracense* | A0A9W4RUJ9 |
| | | | | *Colletotrichum lupini* | A0A9Q8WHQ0 |
| | | | | *Colletotrichum scovillei* | A0A9P7UCT2 |
| | | | | *Colletotrichum karsti* | A0A9P6LJE3 |
| | | | | *Colletotrichum plurivorum* | A0A8H6N6W4 |
| | | | | *Colletotrichum musicola* | A0A8H6MUJ8 |
| | | | | *Colletotrichum gloeosporioides* | A0A8H4FMD8 |
| | | | | *Colletotrichum asianum* | A0A8H3WFM0 |
| | | | | *Colletotrichum tanaceti* | A0A4U6XNA3 |
| | | | | *Colletotrichum higginsianum* | A0A4T0VRV8 |
| | | | | *Colletotrichum trifolii* | A0A4R8RVX5 |
| | | | | *Colletotrichum chlorophyti* | A0A1Q8S4U5 |
| | | | | *Colletotrichum orchidophilum* | A0A1G4BDV4 |
| | | | | *Colletotrichum tofieldiae* | A0A166YUD9 |
| | | | | *Colletotrichum incanum* | A0A166N8W8 |
| | | | | *Colletotrichum salicis* | A0A135UNA5 |
| | | | | *Colletotrichum fioriniae* PJ7 | A0A010RT71 |
| | | Plectosphaerellaceae | Plectosphaerella | *Plectosphaerella plurivora* | A0A9P8V827 |
| | | | | *Plectosphaerella cucumerina* | A0A8K0X9Z6 |
| | | | Verticillium | *Verticillium dahliae* (strain VdLs.17 / ATCC MYA-4575 / FGSC 10137) | G2WX33 |
| | | | | *Verticillium dahliae* | A0A2J8CM44 |
| | | | | *Verticillium longisporum* | A0A0G4MZC7 |

**Table S1.** ***(continued)***

| Class | Order | Family | Genus | Species | Uniprot Accession |
|---|---|---|---|---|---|
| Sordariomycetes | Hypocreales | Bionectriaceae | Clonostachys | *Bionectria ochroleuca* (*Gliocladium roseum*) | A0A8H7NKF0 |
| | | Clavicipitaceae | Claviceps | *Claviceps purpurea* (strain 20.1) | M1WHE9 |
| | | | | *Claviceps pusilla* | A0A9P7NGJ6 |
| | | | | *Claviceps africana* | A0A8K0NIP7 |
| | | | Clavicipitaceae *incertae sedis* (no rank) | *[Torrubiella] hemipterigena* | A0A0A1T7K9 |
| | | | Metarhizium | *Metarhizium rileyi* (strain RCEF 4871) | A0A167GU18 |
| | | | Moelleriella | *Moelleriella libera* RCEF 2490 | A0A168DGN7 |
| | | | Pochonia | *Pochonia chlamydosporia* 170 | A0A179FPA4 |
| | | | Ustilaginoidea | *Ustilaginoidea virens* | A0A8E5MG02 |
| | | Cordycipitaceae | Akanthomyces | *Cordyceps confragosa* (*Lecanicillium lecanii*) | A0A179I7K1 |
| | | | Beauveria | *Beauveria bassiana* | A0A2N6ND37 |
| | | | Cordyceps | *Cordyceps militaris* (strain CM01) | G3J7D9 |
| | | | Niveomyces | *Niveomyces insectorum* RCEF 264 | A0A167TAL4 |
| | | Hypocreales *incertae sedis* (no rank) | Hapsidospora | *Hapsidospora chrysogenum* (strain ATCC 11550 / CBS 779.69 / DSM 880 / IAM 14645 / JCM 23072 / IMI 49137) | A0A086T0I3 |
| | | Nectriaceae | Cylindrodendrum | *Cylindrodendrum hubeiense* | A0A9P5HH11 |
| | | | Fusarium | *Fusarium zealandicum* | A0A8H4XLA5 |
| | | | Ilyonectria | *Ilyonectria* sp. MPI-CAGE-AT-0026 | A0A8K0Q6X4 |
| | | | Neonectria | *Neonectria ditissima* | A0A0P7B837 |
| | | | Thelonectria | *Thelonectria olida* | A0A9P8WJI3 |
| | | Ophiocordycipitaceae | Drechmeria | *Drechmeria coniospora* | A0A151GV38 |
| | | | Hirsutella | *Hirsutella rhossiliensis* | A0A9P8N0V7 |
| | | | | *Hirsutella minnesotensis* 3608 | A0A0F7ZT60 |
| | | | Ophiocordyceps | *Ophiocordyceps sinensis* (strain Co18 / CGMCC 3.14243) | T5A5G2 |
| | | | | *Ophiocordyceps camponoti-floridani* | A0A8H4VH10 |
| | | | | *Ophiocordyceps sinensis* | A0A8H4PST6 |
| | | | | *Ophiocordyceps camponoti-saundersi* (nom. inval.) | A0A369HL98 |
| | | | | *Ophiocordyceps camponoti-leonardi* (nom. inval.) | A0A369GKI2 |
| | | | | *Ophiocordyceps polyrhachis-furcata* BCC 54312 | A0A367LC49 |
| | | | | *Ophiocordyceps camponoti-rufipedis* | A0A2C5Z811 |
| | | | | *Ophiocordyceps australis* | A0A2C5YFX2 |
| | | | | *Ophiocordyceps unilateralis* | A0A2A9PIC9 |
| | | | Purpureocillium | *Purpureocillium takamizusanense* | A0A9Q8V6V0 |
| | | | | *Purpureocillium lilacinum* | A0A179HH70 |
| | | | Tolypocladium | *Tolypocladium capitatum* | A0A2K3QLA3 |
| | | | | *Tolypocladium ophioglossoides* (strain CBS 100239) | A0A0L0N740 |
| | | Stachybotryaceae | Stachybotrys | *Stachybotrys elegans* | A0A8K0SK14 |

**Table S1.** ***(continued)***

| Class | Order | Family | Genus | Species | Uniprot Accession |
|---|---|---|---|---|---|
| Sordariomycetes | Magnaporthales | Magnaporthaceae | Gaeumannomyces | *Gaeumannomyces tritici* (strain R3-111a-1) | J3PEG6 |
| | | | Magnaporthiopsis | *Magnaporthiopsis poae* (strain ATCC 64411 / 73-15) | A0A0C4DQC1 |
| | Ophiostomatales | Ophiostomataceae | Ophiostoma | *Ophiostoma piceae* (strain UAMH 11346) | S3D4R0 |
| | | | Sporothrix | *Sporothrix schenckii* (strain ATCC 58251 / de Perez 2211183) | U7PN22 |
| | Sordariales | Chaetomiaceae | Chaetomium | *Chaetomium* sp. MPI-SDFR-AT-0129 | A0A8K0LP65 |
| | | | Thermochaetoides | *Chaetomium thermophilum* (strain DSM 1495 / CBS 144.50 / IMI 039719) | G0RYY6 |
| | | | Thermothelomyces | *Thermothelomyces thermophilus* (strain ATCC 42464 / BCRC 31852 / DSM 1799) | G2Q5H3 |
| | | Podosporaceae | Podospora | *Podospora anserina* (strain S / ATCC MYA-4624 / DSM 980 / FGSC 10383) | B2ATD0 |
| | | Sordariaceae | Neurospora | *Neurospora crassa* (strain ATCC 24698 / 74-OR23-1A / CBS 708.71 / DSM 1257 / FGSC 987) | V5IQ65 (Trx1A) |
| | | | | *Neurospora crassa* (strain ATCC 24698 / 74-OR23-1A / CBS 708.71 / DSM 1257 / FGSC 987) | V5IQJ8 (Trx1B) |
| | | | | *Neurospora tetrasperma* (strain FGSC 2509 / P0656) | G4U706 |
| | | | Sordaria | *Sordaria macrospora* (strain ATCC MYA-333 / DSM 997 / K(L3346) / K-hell) | F7VQD9 |
| | | Sordariales *incertae sedis* (no rank) | Madurella | *Madurella mycetomatis* | A0A175VYT6 |
| | Sordariomycetes *incertae sedis* (no rank) | Thyridiaceae | Thyridium | *Thyridium curvatum* | A0A507AN44 |
